# Spinal cord structural and functional architecture and its shared organization with the brain across the adult lifespan

**DOI:** 10.1101/2025.10.02.679488

**Authors:** Caroline Landelle, Nawal Kinany, Samuelle St-Onge, Ovidiu Lungu, Dimitri Van De Ville, Bratislav Misic, Véronique Marchand-Pauvert, Benjamin De Leener, Julien Doyon

## Abstract

The spinal cord connects the brain to peripheral systems. Yet its integration with cerebral networks remains a key neuroscience question. Capturing structural and functional central nervous system (CNS) changes throughout the lifespan is essential for characterizing healthy and pathological aging. Leveraging a unique multimodal dataset combining spinal and cerebrospinal imaging, we jointly mapped the spinal cord structural and functional architecture across adulthood. Our results revealed age-related changes across these modalities and identified organizational principles shared with the brain. These changes were most pronounced in the somatosensory pathway, with microstructural decline coupled to shifts in functional connectivity and local spontaneous activity as aging progresses. Extending analyses to the brain uncovered convergent CNS-wide aging mechanisms, including gray matter loss, functional dedifferentiation, and increased spontaneous activity, highlighting shared neural aging trajectories. Together, our findings provide a systems-level view of alterations with age and lay the groundwork for early biomarkers of sensorimotor decline.

## 1. Introduction

Mapping the brain connectome is a core pursuit in neuroscience aimed at understanding how the anatomical architecture shapes functional interactions and how these evolve across the lifespan. Key findings indicate that brain microstructural and functional features follow a sensory-to-associative hierarchy ^1–4^, characterized by stronger coupling in sensory and motor cortices compared to associative areas. This hierarchical organization reflects a functional organization from perception and action to integration ^5^, which evolve across the lifespan and in diseases ^6–8^. However, these investigations are limited to the brain and are overlooking the spinal cord; a critical component of the central nervous system (CNS) that relays and processes somatomotor information between the brain and the periphery. This gap raises important questions: does the spinal cord exhibit properties similar to those of the cortical sensory and motor areas, such as strong similarities among functionally related spinal segments? How does this organization evolve across the lifespan? Mapping the healthy structural and functional architecture of the spinal cord is thus crucial for detecting deviations that may underlie pathological aging and neurological disorders.

Recent neuroimaging advances have provided unique opportunities to assess both functional and structural properties of the spinal cord *in vivo* (Cohen-Adad et al., 2021; Kinany et al., 2022; Landelle et al., 2021), and to image the brain and spinal cord simultaneously ^12,13^. Concomitant advances in spinal cord-specific analysis pipelines have enabled direct functional connectivity mapping between multiple spinal segments in both health and disease ^14–19^ and unveiled the large-scale cerebrospinal topography of the somatomotor pathways *in vivo* ^20–22^. Altogether, these advances have made it possible to deepen our understanding of somatomotor pathway architecture, from the cortex down to the spinal cord.

Importantly, the neural bases that support somatosensory and motor function undergo significant changes across the lifespan resulting in altered sensorimotor performances with aging ^23–25^. Elucidating how aging affects the different CNS structures is thus essential for delineating healthy aging trajectories, identifying periods of heightened vulnerability to sensorimotor disorders such as Parkinson’s disease, and for predicting how aging progresses in individuals with spinal cord conditions. Although substantial research has explored these changes at the brain level (Gooijers et al., 2024; Landelle et al., 2021; Maes et al., 2017) few studies have focused on the spinal cord’s involvement. Notably, existing literature on the aging spinal cord has mainly investigated structure and function separately. Structural investigations have primarily reported age-related demyelination or reductions in cross-sectional area using neuroimaging techniques ^29–32^, while functional changes, such as hyperactivity, have been inferred from electrophysiological studies in non-human models (Iwata et al., 2002; Mayhew et al., 2019; Mayhew et al., 2020). To date, however, no study has directly linked age-related structural and functional changes along the cervical spinal cord *in vivo*, nor examined how these changes reflect those observed at the brain level, leaving a critical gap in our understanding of spinal cord aging and its contribution to sensorimotor decline.

In this study, we acquired a unique and comprehensive adult lifespan neuroimaging dataset (participants from 20 to 80 years) to characterize cervical spinal cord microstructure and simultaneously brain and spinal cord function *in vivo.* Using multimodal spinal MRI, we examine spinal architecture, structure-function coupling, and age-related alterations. We expected that due to the high specificity of sensorimotor processing ^2^, regions within the same spinal segments would share similar functional and structural characteristics. We further hypothesised that aging would be associated with spinal cord microstructural atrophy ^29^ and functional changes ^34,36^, which would manifest differently in the dorsal and ventral pathways due to their distinct roles in somatosensory and motor processing, respectively, as well as their cellular composition. Building on this expectation of region-specific changes and extensive literature on the aging brain ^37^, we also predicted that functional age-related changes in the spinal cord would be linked to structural alterations. Lastly, we conducted structural and functional MRI, simultaneously for the brain and spinal cord, enabling to extend our understanding of the hierarchical organization from the cortex down to the spinal cord, and to investigate the age-related changes across CNS levels. We then hypothesized that structural and functional changes in the spinal cord, including atrophy, functional dedifferentiation, and alterations in local dynamics, parallels those observed in the brain, hence supporting the existence of shared CNS-wide aging trajectories.

## 2. Results

This cross-sectional study included three microstructural spinal cord imaging contrasts: T2*-weighted (T2s), magnetization transfer (MT) and diffusion-weighted imaging (DWI), as well as structural (T1w: T1-weighted) and functional MRI volumes covering simultaneously the brain and cervical spinal cord (from the top of the brain to T1 vertebral level) in 70 healthy right-handed participants aged 20 to 80 years old (36 females, Fig. S2A and Table S1). After quality control, the final sample comprised 67 participants with T2s, T1w, and fMRI imaging, of whom 55 had also MT and/or DWI data. Spinal cord analyses were performed using the Frostell’s segmental level parcellation (Frostell et al., 2016), with each segment subdivided into four parcels: dorsal-right (DR), dorsal-left (DL), ventral-right (VR) and ventral-left (VL, Fig. 1A). To validate the Frostell‘s segmental organization in our dataset, we employed a data-driven approach to extract functional segments (Fig. S3A; Kinany et al., 2024). We found a high degree of overlap between the Frostell’s atlas and the extracted components, as evidenced by a Dice coefficient of 0.80 ± 0.03 (Fig. S3B).

**Figure 1.**
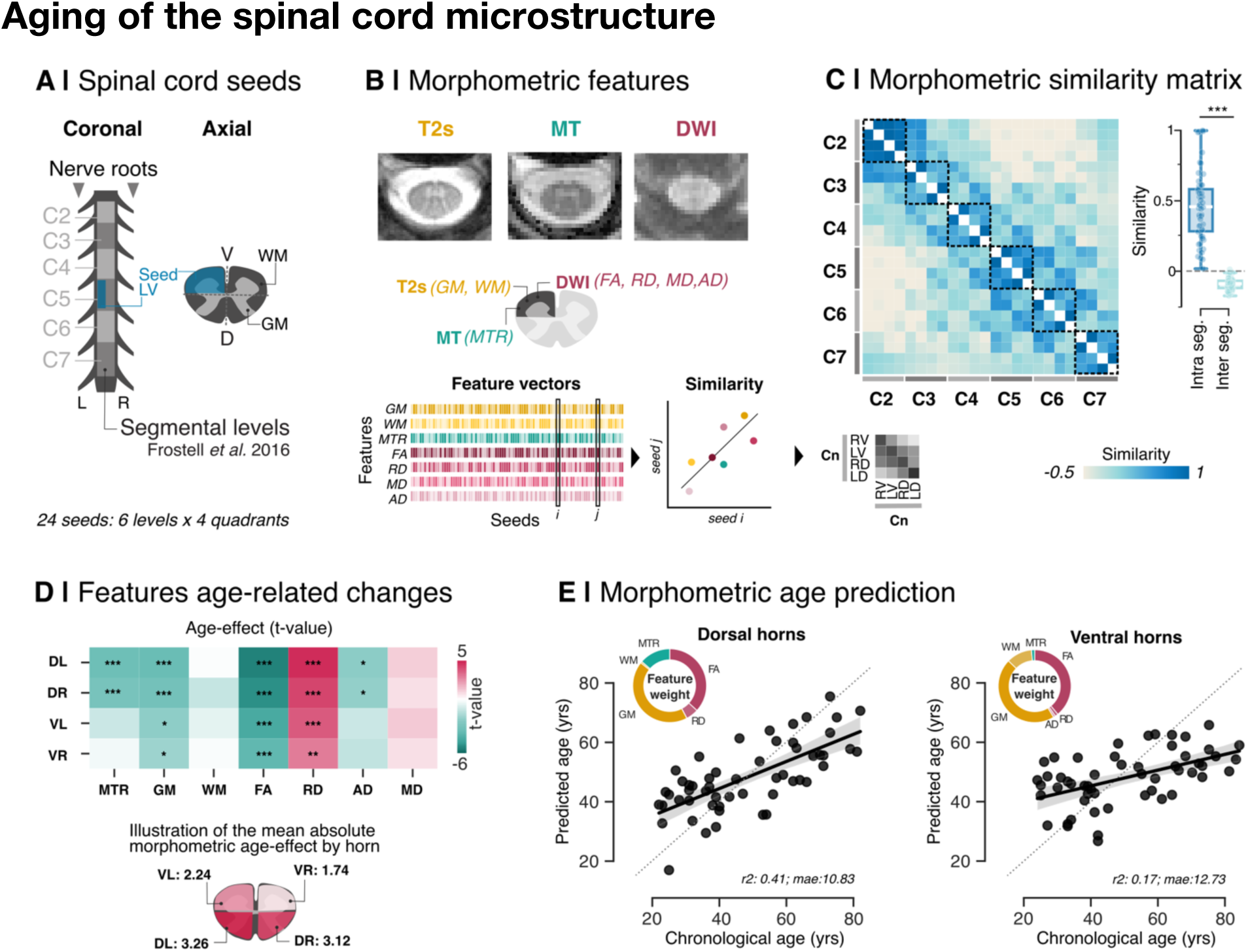
Age related changes of the spinal cord microstructure. **A.** Spinal cord images were parcellated into 24 regions [C2 to C7 Frostell segment subdivided into four horns: dorsal-right (DR), dorsal-left (DL), ventral-right (VR) and ventral-left (VL)]. **B.** We extracted microstructural features from T2s contrast (GM: gray matter voxel counts, WM: white matter voxel counts), the magnetization transfer (MTR: magnetization transfer ratio) and the diffusion (DWI) image (FA: fractional anisotropy, RD: radial diffusivity, MD: mean diffusivity, AD: axial diffusivity). **C.** A morphometric similarity matrix was computed using pairwise Pearson correlation between all possible spinal regions (24×24). Intra-and inter-segment similarity were compared using a two-tailed paired t-test **D.** The heatmap represents t-values reflecting the age-related effect on each microstructural feature (green indicates a decrease with age, pink an increase). The mean absolute effect across features is illustrated on an axial view of the spinal cord for each quadrant. **E**. Age prediction based on the combined microstructural features for dorsal or ventral horn regions. The donut plot illustrates the percentage of contribution (or weight) of each feature to the prediction model.

The results are organized as follows. We first investigated whether different spinal regions exhibit similar microstructural organization and how this organization changes with aging. Second, we studied the spinal cord functional organization and its age-related alterations. Third, we characterized the coupling between spinal structural and functional organization in relation to aging. Finally, we examined whether structural and functional age-related changes in the spinal cord parallel those observed in the brain.

### Aging of the spinal cord microstructure

To characterize accurately the network organization of the spinal cord, we integrated multiple microstructural MRI features that capture distinct anatomical properties across regions. More specifically, we extracted seven microstructural features (Fig. 1B): T2*w-based GM and WM voxel counts, MT ratio (MTR) from MT imaging and diffusion-derived metrics from DWI (*i.e.*, FA: fractional anisotropy, RD: radial diffusivity, MD: mean diffusivity, AD: axial diffusivity) between C2 and C7 segments. We then quantified the morphometric similarity across spinal cord regions by calculating the pairwise correlation between regional feature vectors, comprising the seven microstructural features measured locally for each region and each individual. Interestingly, regions belonging to the same spinal cord segment exhibited greater similarity than those from other cervical segments (Fig. 1C, paired two-tailed t-test; *t*(53)= 20.81, *p*<0.001).

Next, we examined age-related changes in the microstructural organization of each spinal cord parcels to determine whether aging affects the spinal cord uniformly or follows a region-specific pattern. Using an ordinary least squares (OLS) model including age and sex as fixed effects, we estimated the linear age-related alterations for each feature across the four spinal quadrants (*i.e.*, DR, DL, VR, VL). Significant age-related decreases were observed for MTR, GM voxels count, FA, and AD, whereas RD showed an increase (Fig. 1D, and Table S2). The strongest changes were found in the dorsal-left (DL) horn (MTR: p_FDR_<0.001, GM: p_FDR_<0.001, FA: p_FDR_<0.001, RD: p_FDR_<0.001, AD: p_FDR_=0.017). Averaging the absolute t-values across features confirmed that microstructural changes were more pronounced in the DL horn (*t*= 3.26 ± 1.99), followed by DR (*t*= 3.12 ± 1.64), VL (*t*= 2.24 ± 1.52) and VR (*t*=1.74 ± 1.44) horns. To further visualize age-related patterns, we generated average metric maps for different age subgroups (Fig. S4). Note that sex effects are also reported in supplementary materials (Table S2) ans show increase in GM voxels count in males comapred to females and also sex-effects on d. To confirm whether the spinal cord microstructural aging follows a region-specific, distinctly affecting the dorsal (somatosensory pathway) and ventral hemi-cord (motor pathway), we employed two elastic net regression models incorporating all extracted features to predict participants’ age. Microstructural features predicted age more accurately in the dorsal spinal cord (Mean absolute error, MAE: 10.83 years, r^2^=0.41) than in the ventral regions (MAE: 12.73 years, r^2^=0.17, Fig. 1E). In both models, GM voxel counts and FA contributed most to the predictions (dorsal GM counts: 43% of the explained variance, FA: 37%, MTR: 13.61%, RD: 5.56%, WM counts: 0.73%, AD: 0%, MD: 0%WM: 11.38%; ventral GM counts: 46%, FA: 39%, WM counts: 11.37%, MTR: 1.39%, RD: 1.37%, AD: 0.88%,MD: 0%). Lastly, complementary analyses were performed within the three white matter tracts (dorsal column, lateral funiculi and ventral funiculi) to assess tract-specific microstructural changes in the spinal cord. Results showed that the dorsal column (proprioceptive and tactile fibers) exhibited the strongest age-related effect in MTR, FA and RD (see details in Fig. S5). Lower but significant age-related changes were also observed for the three metrics in the lateral funiculi (mainly motor fibers), whereas in the ventral funiculi (nociceptive and thermoceptive fibers), only FA and RD showed significant changes with aging. Overall, these findings suggest that aging of the spinal cord microstructure is more pronounced in somatosensory ascending pathways than in motor descending pathways.

### Aging effects on the functional topography of the spinal cord

To extend our analysis beyond morphometric organization, we characterized the functional architecture of the spinal cord by evaluating distinct functional properties across its regions. As a first step in characterizing spinal cord functional organization, we employed the most widely used fMRI approach for assessing spinal cord function at rest: inter-regional functional connectivity (SpiFC, Eippert et al., 2017; Kaptan et al., 2023; Kong et al., 2014; Landelle et al., 2023). This approach estimates the degree of synchronized activity across spinal cord regions by computing pariwise correlations between their timeseries. Based on the correlation coefficient, we constructed SpiFC matrix across 28 spinal regions (C1 to C7 segments, each divided into four quadrants, Fig. 2b). We then mapped intra-regional dynamic profiles of spinal cord activity (SpiDyn) by analyzing the temporal characteristics of timesries fluctuations within each region. This second approach aimed to capture the dynamical signature of spinal cord function, offering complementary insight beyond static connectivity measures. Specifically, SpiDyn included the amplitude of low fluctuation frequency (ALFF); a commonly used fMRI biomarker of local spinal activity ^41–43^ alongside a set of timeseries features derived from the catch-22 toolbox (Lubba et al., 2019), which quantify the dynamic properties of the spontaneous fMRI signals such as signal complexity, temporal predictability and variability.

**Figure 2.**
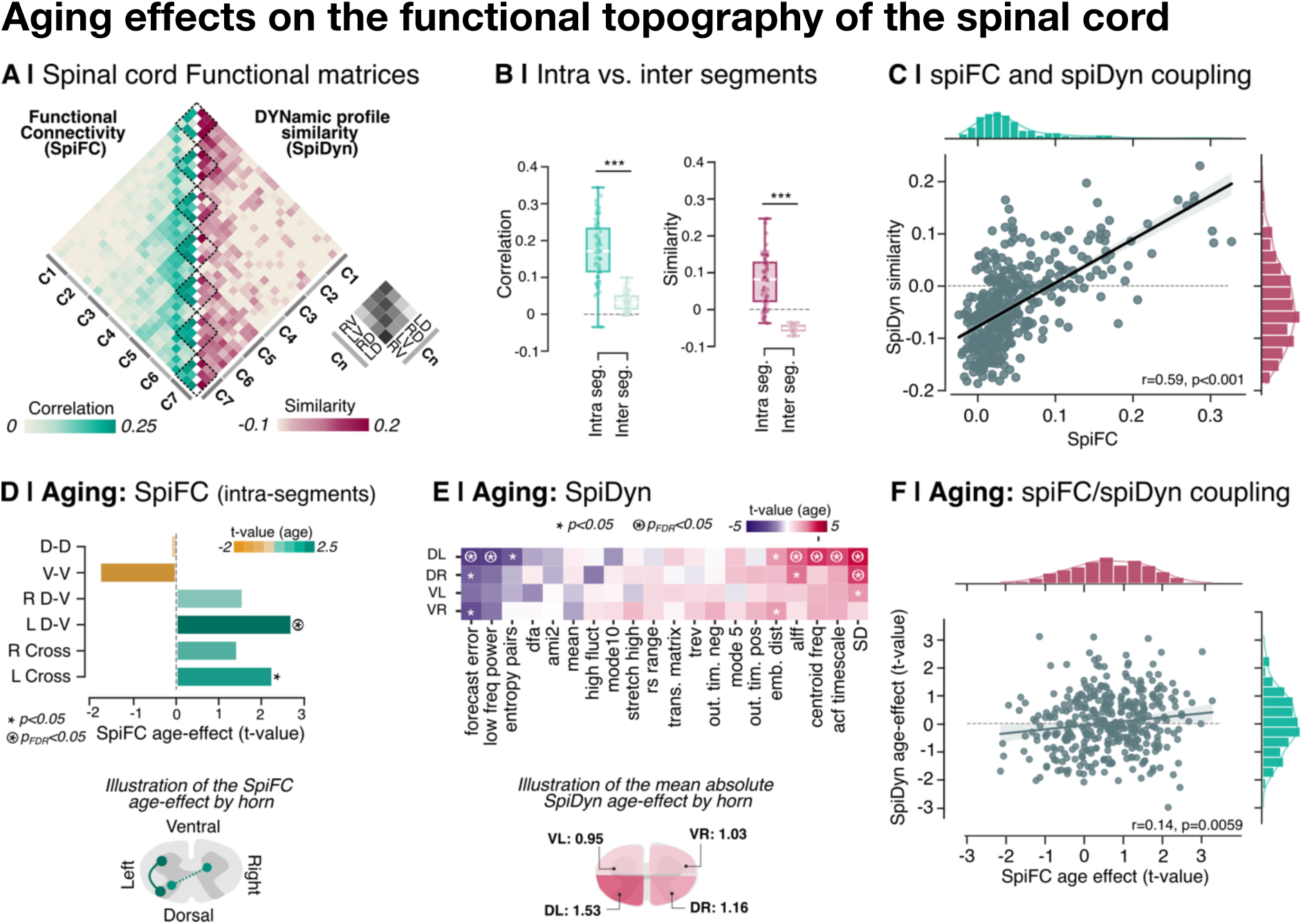
Age-related changes of the spinal cord functional topography. **A.** Spinal cord functional connectivity (SpiFC, green) matrix and dynamic profile similarity (SpiDyn, pink) matrix were computed using pairwise Pearson correlation between all possible spinal regions (28×28). **B.** Intra-and inter-segment SpiFC (green) and SpiDyn (pink) were compared using a two-tailed paired t-test. Both were significantly higher for regions belonging to the same segment (***: p<0.001) **C.** SpiDyn was positively correlated with SpiFC (r denote the Spearman coefficient of correlation). Each dot represents a pairwise combination of spinal regions, and a linear regression line was overlaid on the scatter plot for visualization. **D.** Bar plots show t-values reflecting age-related effects on intra-segmental SpiFC for each of the six types of connexions (D-D: dorso-dorsal, V-V: ventro-ventral, R D-V: right dorso-ventral, L D-V: left dorso-ventral, R Cross: right cross-horns, L Cross: left cross-horns). Increased t-values are shown in green, while decreased are shown in yellow. Functional connectivity increases for L D-V and L Cross connections were significant (*: p=0.0065 and p=0.023), but only L D-V remained significant after FDR correction (circled asterisk, pFDR=0.039). **E.** Heatmap of t-values reflecting age-related effects on each SpiDyn feature (violate indicates a decrease with age, pink an increase). Significant results before and after FDR correction are marked with an asterisk or a circled asterisk, respectively. The mean absolute effect across features is illustrated on an axial view of the spinal cord for each quadrant. **F.** Correlation between aging effects on SpiDyn and SpiFC at each spinal region. We found a significant positive correlation between these two functional metrics related to aging (r denote the Spearman coefficient of correlation). Each dot represents a pairwise combination of spinal regions, and a linear regression line was overlaid on the scatter plot for visualization. Cn: cervical spinal cord level. DL: dorsal left, DR: dorsal right, VL: ventral left, VR: ventral right.

Similar to the morphometric similarity approach, we quantified SpiDyn similarity across the 28 spinal parcels by computing pairwise correlations between regional feature vectors (Fig. 2b). Results revealed that spinal regions within the same spinal segment had significantly greater functional connectivity and dynamic profile similarity than those from different segments (paired two-tailed t-test for SpiFC: t(66)= 17.25, p<0.001 and SpiDyn: t(66)= 15.52, p<0.001). We also found a positive correlation between SpiDyn and SpiFC (Spearman’s r=0.59, p<0.001), hence suggesting that spinal regions with similar local dynamics exhibit synchronized spontaneous activity.

To investigate whether aging impacts this spinal cord functional organization, we first estimated age-related changes in SpiFC. The results revealed a significant increase in average inter-segmental connectivity with age (t(65)=2.93, p=0.0033), whereas intra-segmental connectivity showed no significant change (t(65)=085, p=0.39). When examining the six specific connection types (*i.e.*, dorso-dorsal, ventro-ventral, left dorso-ventral, right dorso-ventral, left cross-horns, right cross-horns), inter-segmental connectivity significantly increased with age across all connections except the dorso-dorsal (Table S3). For intra-segmental connectivity, age-related increases were observed in the left dorso-ventral (t= 2.71, p= 0.0065, p_FDR_= 0.039) and left cross-horns connections (t= 2.26, p= 0.023, p_FDR_= 0.069), although the latter did not survive correction for multiple comparisons (Fig. 2D and Table S4). The increase in inter-segmental connections suggests a decline in the specificity of spinal connectivity with age, alongside selective increases in intra-segmental FC, particularly those driven by the left dorsal horn.

We further examined age-related changes in local spinal cord dynamics (Fig. 2E). The left dorsal horn showed the most pronounced changes, with several SpiDyn features exhibiting a significant increase with aging after multiple correction (standard deviation, autocorrelation timescale, centroid frequency and ALFF) or decrease (Forecast error, low-frequency power) (Table S5). Standard deviation also increased in the right dorsal horn. These patterns suggest that dorsal horns, especially the left side, exhibit age-related changes characterized by increased signal variability and amplitude (*i.e.*, higher standard deviation and ALFF), alongside with reduced signal complexity (*i.e.*, lower low-frequency power) and enhanced predictability (*i.e.*, longer autocorrelation timescale and centroid frequency, decreased forecast error).

Finally, we investigated local dynamic features that showed statistically significant age-related changes to investigate whether alterations in inter-regional spiFC, measured independently from SpiDyn, were associated with local dynamic properties evolving with age across spinal regions. Specifically, we quantified the age-related effect for each pairwise connection in the SpiFC matrix and each pairwise similarity in the SpiDyn matrix before assessing their relationship using Spearman correlation. Notably, the results revealed a weak but significant positive correlation (r=0.15, p=0.003), indicating that spinal regions exhibiting age-related changes in functional connectivity also show corresponding age effects in their time-series dynamic similarity profiles. These results suggest a simultaneous alteration in both spontaneous inter-regional connectivity and local intrinsic dynamics with aging at the spinal cord level. Recognizing the weak association is important as it may reflect a combination of shared and distinct neurobiological processes, highlighting the need to assess both functional aspects to comprehensively capture age-related changes. Sex effects were systematically tested and are reported in supplementary material (Tables S3-S5). We observed sex differences in SpiFC, with females showing increased intra-segmental connectivity compared to males, whereas SpiDyn analyses revealed higher ALFF values in males than females in the dorsal hemicord. These findings highlight the importance of accounting for sex as a variable in the analysis.

#### Aging of the spinal cord structure-function coupling

We next aimed to determine whether functional connectivity (SpiFC) and regional activity patterns (SpiDyn) mirror the microstructural architecture of the spinal cord. The results revealed a strong and significant correlation between morphometric similarity and both SpiFC (Spearman’s r=0.60, p<0.006, Fig. 3A-B) and SpiDyn similarity measures (Spearman’s r=0.66, p<0.006, Fig. 3D-E). These findings suggest that the spinal cord organization, encompassing both inter-regional connectivity and local dynamics, may be shaped by its microstructural architecture.

**Figure 3.**
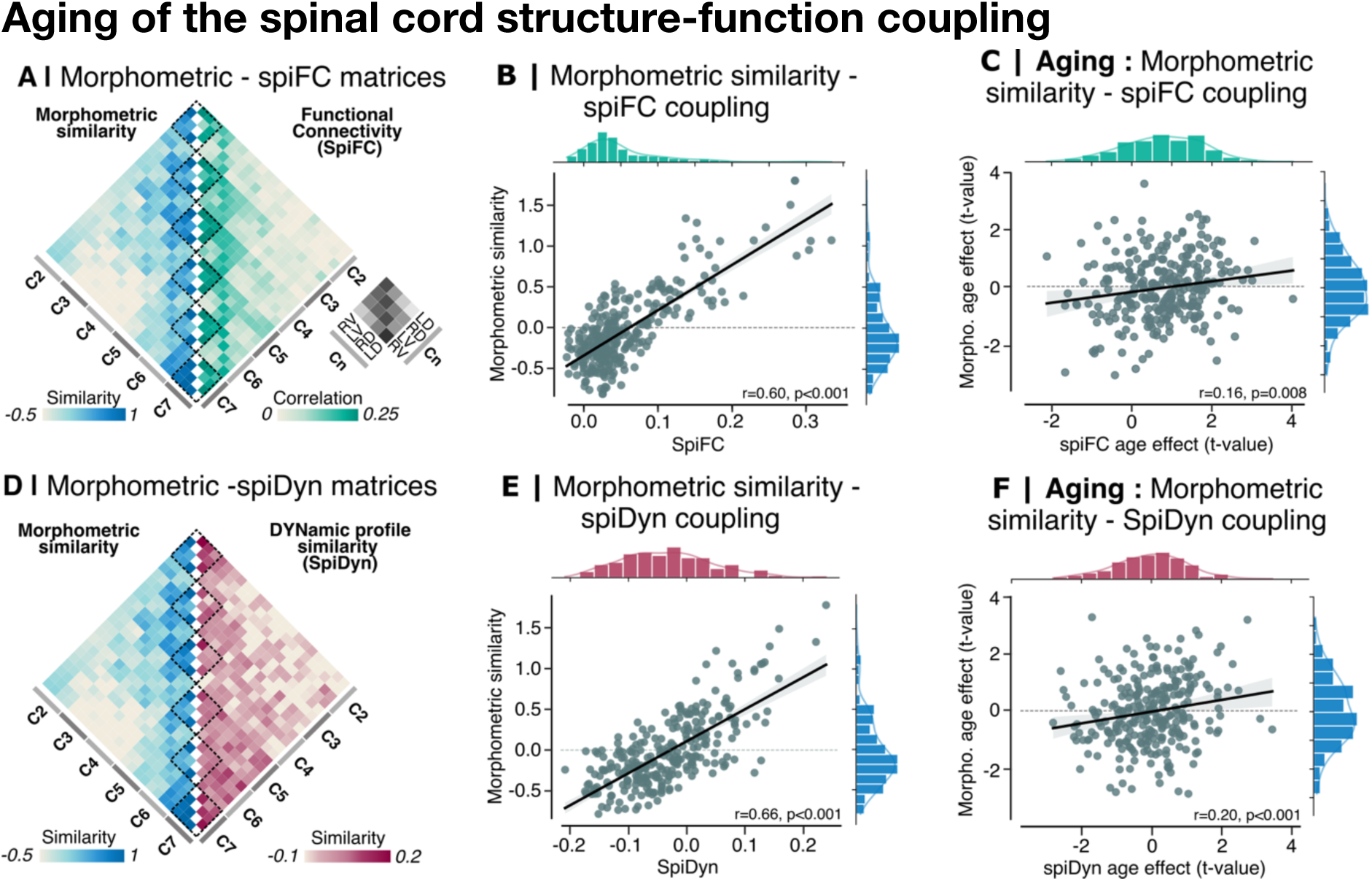
Age-related changes in spinal cord structure-function coupling. A, D. Spinal cord morphometric similarity matrices (blue, A and D) and spinal cord functional connectivity (SpiFC) matrix (green, A) or spinal cord dynamic profile similarity (pink, D) were computed using pairwise Pearson correlation between all possible spinal regions (24×24). B, E. The couplings between morphometric similarity and SpiFC (B) or SpiDyn (E) were assessed using Spearman correlation. We found a strong positive correlation between the two matrices. C, F. Age-related effects on morphometric-SpiFC (C) or morphometric-SpiDyn (F) coupling were assessed by correlating the age effects of morphometric similarity and SpiFC or SpiDyn for each pairwise combination of spinal regions. For plots B, C, E, and F, each dot represents a pairwise combination of spinal regions, a linear regression line was overlaid on the scatter plot for visualization, and r denotes the Spearman coefficient of correlation.

Next, to assess whether aging similarly affects both functional and structural organization of the spinal cord, we estimated how the strength of coupling between microstructural and functional measures (*i.e.*, SpiFC and SpiDyn) is modulated by age. Correlation analyses revealed that microstructural-functional coupling increases with aging, both for inter-regional functional connectivity (Spearman’s r: 0.16, p=0.008, Fig. 3C) and intra-regional dynamic profiles (Spearman’s r: 0.20, p<0.001, Fig. 3D).

Altogether, these results highlight a strong coupling between microstructural and functional organization at the spinal cord level, with structural age-related effects being linked to changes in functional connectivity and local dynamics.

### Parallel structural and functional age-related changes of the brain and spinal cord

To investigate the structural and functional hierarchical organization from the cortex down to the spinal cord and its age-related changes across CNS levels, we conducted structural and functional MRI scans, the latter being acquired simultaneous for the brain and spinal cord. First the relationship between spinal cord and brain structural integrity was assessed to determine whether age-related changes occur in parallel across regions. We performed brain-level analyses across 200 brain cortical regions, 24 cerebellar regions and six subcortical structures (combining Schaefer and Cobra atlases, Schaefer et al., 2018; Tullo et al., 2018). This parcellation enabled the investigation of nine brain networks, along with the whole cervical spinal cord network. To simplify the description of the results, in the main text, we focused on the networks related to somatomotor processing (*i.e.*, the somatomotor, spinal cord, cerebellar and subcortical networks) in Fig. 4A-B, while the results from the other networks are provided in the Supplementary Material. Across these three brain networks, we did not find any significant relationship between the spinal cord GM cross-sectional area (CSA) and the brain GM volume (Fig. 4A, Subcortical: t(66)= -0.62, p=0.53; Cerebellum: t= -1.31, p=0.19; Somatomotor: t(66)= -1.73, p=0.088). Furthermore, there were no significant differences in the age effect between the spinal cord CSA and the subcortical or somatomotor region volume. However, the spinal cord appeared to be more strongly affected by age compared to the cerebellum, as indicated by a significant age-by-region interaction (t=2.19, p=0.031). Importantly, all ten networks revealed strong volumes or CSA decreasing with aging (Fig. 4B, Fig. S6 and Table S6).

**Figure 4.**
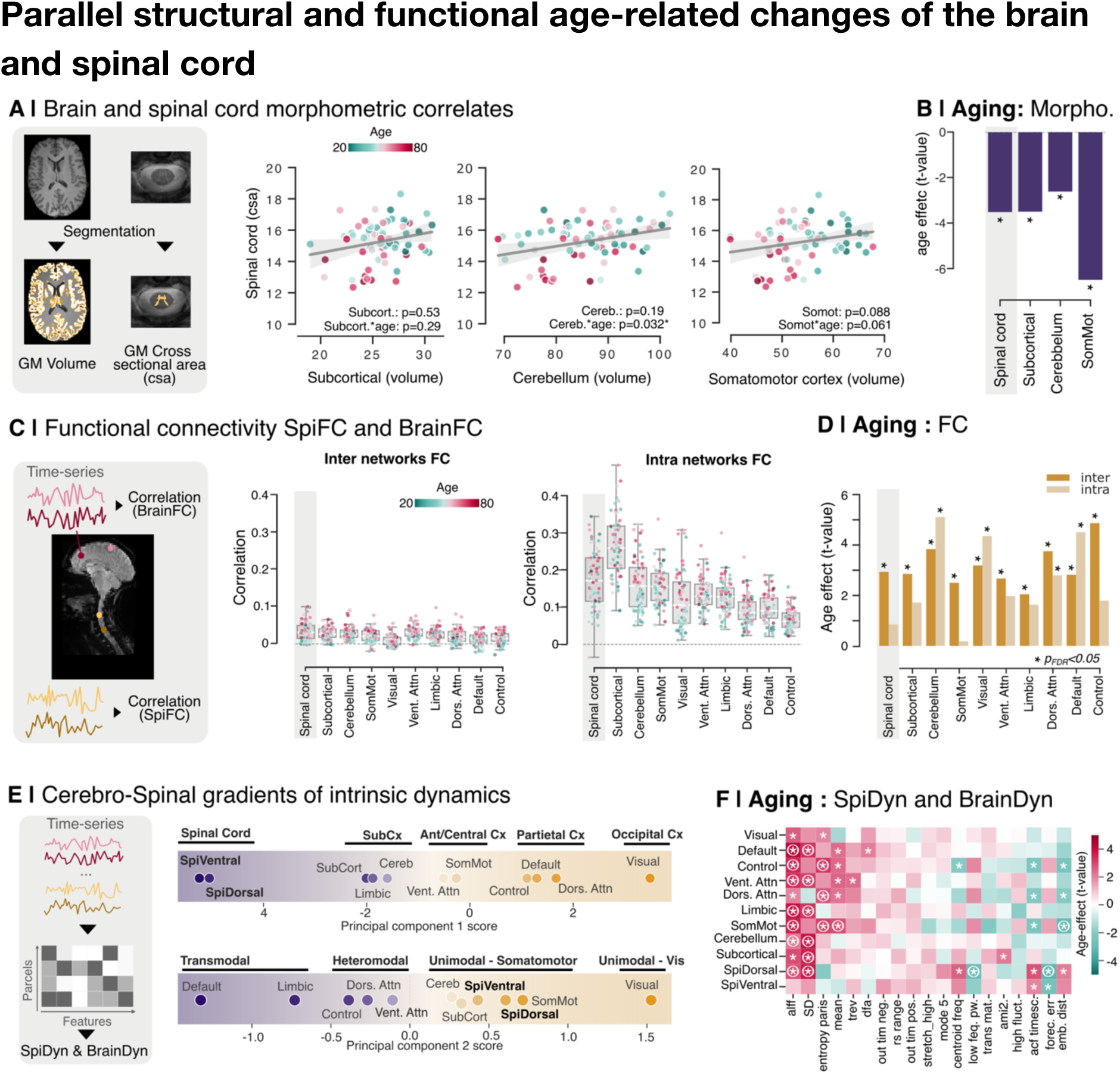
Brain and spinal cord correlates and their age-related changes. **A.** Cerebral gray matter (GM) volume (in mL) and spinal cross-sectional area (CSA, in mm^3^) were obtained after segmentation of the brain T1w contrast and the spinal cord T2s contrast respectively. Linear regression between CSA and brain GM volume are shown for subcortical area, cerebellum and somatomotor cortex. Reported p-values correspond to the main effect of brain area and the brain area*age interaction. Each dot represents an individual participant (n=67), color-coded by age from green (younger) to pink (older). **B.** Spinal cord GM CSA and GM volume for each brain network were entered in a regression model including age and sex as fixed effects. Bar plots represent age-related effects (t-values) for each network (spinal cord network highlighted in gray). Asterisks indicate statistically significant effect after FDR correction. **C.** Brain and spinal cord time-series were extracted and correlated to measure brain and spinal cord functional connectivity, respectively. Boxplots show intra-(left) and inter-networks (right) FC across the ten networks, with the spinal cord network highlighted in gray. Each dot is an individual participant (n=67), color-coded by age from green (younger) to pink (older). **D.** FC values were entered in a regression model including age and sex as fixed effects. Bar plots represent age-related effects (t-values) for each network (spinal cord network highlighted in gray), with yellow representing inter-and pale-yellow intra-network FC. Asterisks indicate statistically significant effect after FDR correction. **E.** Parcel-specific intrinsic dynamic features were derived from time-series across both brain and spinal cord structures. Next, principal component analysis (PCA) was applied to the parcels x time-series features matrix to identify dominant patterns of time-series dynamic variation. To access how strongly each network expresses the first (top) or second (bottom) principal component, we projected the feature matrix onto the PCA-derived eigenvectors, yielding a score for each parcel. Each dot represents the centroid score of parcels within a network, positioned along the gradient defined by the respective PCA score. **F.** Heatmap representing t-values for age-related effects on each dynamic feature across networks (green: decrease with age, pink: increase). Significant results before and after FDR correction are represented with an asterisk or a circled asterisk, respectively.

Next, we aimed to deepen our understanding of the functional organization spanning the brain and the spinal cord by computing the average intra-network FC (*i.e.*, FC between parcels within the same network) and inter-network FC (*i.e.*, FC between parcels belonging to distinct networks) for each of the ten networks. Across all networks, intra-network FC was consistently higher than inter-network FC (Fig. 4C and Table S7; t(66)= 40, p<0.001), while a weaker but positive FC was observed between networks at each CNS level (t(66)= 14.8, p<0.001; Table S8). This pattern held for both brain and spinal cord networks, suggesting a shared organizational principle of network segregation at rest, characterized by strong coherent FC within the same network and relatively weak but consistent FC across them. Next, we examine age-related functional connectivity changes in both brain and spinal cord networks to determine whether functional reorganization with aging follows a system-wide pattern or emerges differentially across specific networks. The aging effect was assessed using OLS regression model for both intra-and inter-network FC, including age and sex as fixed effects. We found a significant increase in the spinal cord and brain inter-network FC with aging, suggesting a shift toward less specific FC patterns, potentially reflecting dedifferentiation processes ^47^ at each CNS level (Fig. 4D, F Table S9). In contrast, spinal cord intra-network FC did not exhibit significant age-related alterations, consistent with most brain networks, except for the posterior networks (visual, dorsal attentional and default mode) and the cerebellum, which showed an increase in intra-network FC with aging (Fig. 4D, Table S10).

Finally, we investigated the topographical organization of time-series dynamics across each level of the CNS, aiming to uncover a comprehensive functional organization spanning the spinal cord, subcortical structures and cerebral cortex. Principal component analysis (PCA) applied to the parcels-by-features matrix extracted independent patterns of intrinsic dynamics, with the first two components collectively accounting for 62% of the variance. The first principal component (PC1) captured a cortical-subcortical-spinal cord gradient of intrinsic dynamics (Fig. 4E, top). Within the cortex, this gradient followed the well-established dorsolateral-ventromedial axis of functional organization ^48,49^. The second component (PC2) revealed a unimodal-transmodal gradient of intrinsic dynamics (Bernhardt et al., 2022; Huntenburg et al., 2018; Margulies et al., 2016; Shafiei et al., 2020; Fig. 4E, bottom). The unimodal end of this gradient comprised the visual cortex, followed by somatomotor regions, including the somatomotor cortex, spinal cord and subcortical structures (cerebellum, basal ganglia). Progressing along the gradient, it encompassed heteromodal associative areas capable of integrating information across modalities and finished in default mode regions, which are widely recognized as transmodal. To assess age-related changes in dynamic functional organization across the CNS, we applied OLS regression to each time-series feature, including age and sex as fixed effects. Notably, ALFF and standard deviation (SD), two key metrics of signal amplitude fluctuation, significantly increased with age across several networks (Table S11). These widespread changes suggest that aging is associated with a general increase in spontaneous neural fluctuation throughout the CNS.

## 3. Discussion

Despite the spinal cord’s key role in somatomotor function and its clinical relevance in conditions such as spinal cord injury, multiple sclerosis, and age-related pathologies like Parkinson’s disease, *in vivo* neuroimaging of the human spinal cord has historically lagged that of the brain. This disparity stems from significant technical challenges in imaging a deep, narrow structure surrounded by different tissues, cerebrospinal fluid, and moving organs. Over the past decade, however, substantial progress in both acquisition and analysis techniques has enabled more precise microstructural imaging and the extraction of robust functional signals from the spinal cord (Cohen-Adad et al., 2021; Kinany et al., 2022; Landelle et al., 2021). Leveraging these advances, we built a unique multimodal dataset spanning the entire cervical spinal cord and brain in a cross-sectional cohort of 70 healthy young and older adults. This integrative approach allows us to reveal that spinal somatosensory regions (dorsal) are predominantly affected by aging. This approach allowed us to identify region-specific aging effects, quantify structure-function coupling in the spinal cord, and reveal shared organizational principles across the CNS.

We observed pronounced age-related structural and functional changes in the dorsal spinal cord, which is primarily responsible for somatosensory processing. Older adults exhibited reductions in gray matter volume, magnetization transfer ratio (MTR), and diffusion metrics such as fractional anisotropy (FA) and axial diffusivity (AD), alongside increased radial diffusivity (RD), consistent with declines in tissue integrity, potentially reflecting neural shrinkage, impaired axonal organization, or dys/demyelination. These results align with previous research showing that aging leads to a preferential loss of large myelinated sensory fibers, reduced conduction velocity, and impaired somatosensory acuity ^50^. Electrophysiological assessments of proprioceptive-motor loops (H-reflex) in humans have similarly demonstrated greater deterioration in the sensory component compared to the motor component with aging ^51–54^. Extending these findings, our complementary tract-specific microstructural analyses revealed significant demyelination in dorsal WM columns, associated with tactile and proprioceptive afferents, whereas no such changes were observed in WM ventral columns, which mainly contain nociceptive and thermoceptive fibers^55^. This pattern mirrors prior *ex-vivo* study showing that the dorsal column is particularly susceptible to age-related degeneration ^56^, supporting the hypothesis that aging may differentially affect the somatosensory system, with proprioceptive afferents being particularly vulnerable ^23,27,57,58^. Our functional analyses further mirrored these changes: the dorsal cord exhibited increased local spontaneous activity (higher SD and ALFF), reduced signal richness (decreased low-frequency power), and enhanced predictability (increased autocorrelation), along with increased connectivity primarily involving the left dorsal quadrant (*i.e.,* left dorso-ventral and left dorsal-right ventral FC). These findings align with rodent studies showing age-related dorsal hyperexcitability and shifts in excitation-inhibition balance ^33,34,36,59^. Interestingly, in our study, the left dorsal quadrant, which receives somatosensory input from the participants’ non-dominant upper limb, was most affected, suggesting that reduced limb use may accelerate aging in its corresponding spinal territory —a potential target for somatosensory rehabilitation.

The literature on age-related changes in the ventral horn is inconsistent regarding whether motor neurons are lost or undergo atrophy with aging ^60–63^. Nonetheless, there is some evidence of age-related alterations, including a reduction in the excitatory-inhibitory synaptic ratio and an increase in senescent motor neurons ^60,64^. Although the functional effects in the ventral quadrants were less pronounced in our analyses, these previous findings align with the microstructural changes that we also observed in the ventral horns (*i.e.*, reduction in spinal cord gray matter and microstructural integrity with aging). To better understand these region-specific changes with aging along the sensory-motor axis, future studies should investigate spinal cord function beyond resting-state assessments by incorporating motor task or somatosensory-stimuli paradigms, allowing a more comprehensive evaluation of somatosensory and motor alterations associated with aging.

Beyond separate structural and functional analyses, we examined their interplay and how this coupling evolves with aging. We uncovered a close structure-function correspondence along the cervical spinal cord, consistent across measures of functional organization, whether assessed through inter-regional connectivity or local spontaneous activity. This tight coupling mirrors the structural-functional coupling observed in the primary somatomotor cortex, where highly specialized afference and efference support greater correspondence between structure and function compared to other brain regions ^2,48^. Like the somatomotor cortex, the spinal cord exhibited a highly specialized functional architecture, exemplified by reflex loops, the most stereotyped stimulus-response correspondence, which can involve as few as a single synapse between somatosensory afference and motor efference, directly linking sensation and action ^65^. Importantly, the spinal structural-functional coupling is not static but changes with aging, with structural decline associated with altered functional connectivity and local dynamics–the latter showing a stronger association. These findings highlight the need for multidimensional functional metrics to fully capture its reorganization and develop sensitive biomarkers of spinal cord aging or dysfunction. We speculate that the structural-functional coupling changes with aging may reflect neural plasticity, such as the emergence of compensatory mechanisms, but could also result from structural alteration, such as demyelination, that slows and reduces the fidelity of the signal transmission, ultimately leading to functional communication dedifferentiation (*i.e*., stronger inter-segmental connectivity) or greater signal variability (*i.e.,* increase ALFF, standard deviation). Combining behavioral assessments with multimodal imaging will be critical to distinguish adaptive and maladaptive changes and to determine how this structural-functional correspondence evolves in other contexts, such as motor learning, injury recovery or neurodegenerative diseases.

We next asked whether shared structural and functional organizational principles span the architecture of the CNS across its different levels. Mapping the connectome beyond the cortex, particularly in structures like the spinal cord, is essential to understand how the CNS interfaces with the body and the environment as well as how this relationship evolves across the lifespan. Recent studies have begun to integrate multimodal imaging of both cortical and extracortical structures, including the brainstem or the spinal cord, as well as peripheral organ systems, to provide more comprehensive view of CNS-body communications ^20,66,67^. Building on this emerging framework, we leveraged neuroimaging data from simultaneous imaging of the spinal cord and brain within the same individuals, uniquely positioning us to examine CNS-wide patterns of organization and their age-related changes. This approach not only deepened our understanding of brain organization but also enabled us to validate and contextualize our spinal cord findings, revealing common principles of neural architecture and shared age-related changes across the neuroaxis. Notably, shared organizational principles were more apparent in functional than in structural analyses. While functional metrics revealed consistent patterns across spinal cord and brain regions involved in sensorimotor processing, structural volumetric analyses did not show a significant correlation between the spinal cord and brain regions, consistent with previous reports that brain GM volume does not predict spinal cord GM CSA ^31,68^. Examining functional connectivity organization revealed similar patterns of connectivity in both structures, characterized by stronger connectivity within networks and weaker connectivity between distinct networks. Analogous to the brain, inter-network FC in the spinal cord increased with aging. This phenomenon, well-documented in brain regions such as the primary somatomotor cortex ^69–71^, is associated with dedifferentiation (*i.e.,* a decrease in neural specialization), which manifests as more widespread connectivity across networks or a recruitment of additional brain areas during motor tasks and somatosensory stimulation ^23–25^. Our results provide the first evidence that dedifferentiation extends across the CNS.

We further examine local time-series dynamics using a principal functional gradient approach, identifying two continuous, macroscale patterns of intrinsic CNS organization ^1^. One pattern captured a cortical-subcortical-spinal cord axis, spanning the occipital cortex to the spinal cord. This gradient extended the well-established anteroposterior cortical gradient ^48,49^ by integrating subcortical regions and the spinal cord, for the first time, and thereby capturing the neuroaxis of the human CNS along a rostro-caudal coordinate system. The second pattern followed a unimodal-transmodal gradient, with from the visual network to the default mode network ^1,5,48,49^. Remarkably, the cerebellum, subcortical nuclei and the spinal cord were positioned close to the somatomotor network along this gradient, likely reflecting their shared functional involvement in somatomotor processing. Together, these gradients provide compelling evidence for a hierarchical and continuous functional organization across the CNS, bridging cortical, subcortical and spinal levels. We further examined age-related changes in local spontaneous activity throughout the CNS and found that increased signal variation (higher ALFF and standard deviations) was a consistent feature. This pattern suggests that aging is associated with a generalized shift toward less stable neural activity, potentially driven by reduced neural specialization, compensatory recruitment of additional neural resources, or greater physiological noise. These findings reinforce the notion that aging induces widespread functional reorganization, not only in the brain but across the entire CNS.

In the present study, we extended the concept of the *in vivo* cortical connectome to the spinal cord and characterized its age-related changes using multimodal imaging and multidimensional analyses. This opens new avenues for understanding the trajectory of the somatomotor system across the lifespan and holds promise for clinical application. For example, both the brain and spinal cord are affected in Parkinson’s disease ^18,72–74^, yet they are not studied together. Applying integrated structural and functional analyses across these structures could yield deeper insights into disease mechanisms and help identify early biomarkers of somatomotor dysfunctions. Moreover, our observation of age-related region-specific alterations in the spinal somatosensory pathways raises important questions about the underlying peripheral *versus* central mechanisms driving these changes. Future longitudinal studies, combining neuroimaging, electrophysiology and behavioral measurement, will be essential to track the temporal evolution of the CNS-wide reorganization with aging. Ultimately, this work lays the foundation for a more comprehensive understanding of the neural basis of sensorimotor aging and its disruption in neurological conditions.

Despite the strengths of our multimodal and multilevel approach, several limitations warrant consideration. First, although we observed consistent patterns of age-related change, our cross-sectional design limits causal inference regarding the temporal dynamics of CNS aging. Longitudinal studies will be essential to disentangle progressive decline from compensatory reorganization. Second, our study focused on the resting state to characterize functional organization and its age-related changes. It is important to recognize the complementary insights offered by task-based paradigms that can more directly probe functional specialization and recruitment patterns during sensorimotor engagement. Finally, a further limitation of our study is the lack of microstructural imaging data at the brain level, which prevented direct comparisons between spinal and cerebral fine-scale tissue integrity. Future studies incorporating advanced brain microstructural imaging techniques will thus be essential to fully characterize the interplay between structure and function across the entire CNS.

In summary, we leveraged recent advances in neuroimaging to extend the scope of aging research beyond the brain, encompassing the spinal cord. Our findings highlight the continuity and specialization of sensorimotor systems across the neuroaxis and reveal shared patterns of age-related reorganization. This work opens new avenues for identifying CNS-wide biomarkers and therapeutic targets for age-related neurodegenerative disorders.

## 4. Methods

### Participants & MRI acquisitions

Seventy right-handed healthy participants (36 females; age 48.82 ± 16.94 years old, [20-80] years) were included in this study. The experiment was approved by the local ethics committee (MUCH REB 2019-4626), and all participants gave their written consent in accordance with the Helsinki Declaration. The MRI data were acquired at the Neuro (Montreal Neurological Institute, Canada) using a 3-Tesla MRI Scanner (Magnetom-Prisma^fit^, Siemens, Erlangen, Germany) equipped with a 64-channel head and neck coil. Participants were positioned head-first supine and were instructed to relax, minimize motion and swallowing. Throughout the scanning session, the participants wore Neck and Brachial Plexus SatPads, around their neck and on the chest, respectively, as well as physiological sensors that included a pulse sensor on their index and a respiration belt (Siemens Physiology Monitoring Unit).

A high-resolution T1w anatomical image covering the whole brain and the cervical spinal cord up to the first thoracic vertebrae was acquired to facilitate tissue segmentation, as well as vertebral labelling for normalization to the PAM50 template ^75^ for spinal functional and microstructural. This acquisition was followed by simultaneous brain/spinal cord echo planar imaging (EPI) (same field of view as the T1w image) to investigate brain and spinal cord activity during rest (*i.e.,* no explicit task), while participants instructed to refrain from specific thoughts and passively view the *Inscapes* video ^76^. The microstructural images were acquired at the cervical spinal level (C1 to C7 vertebrae), including T2*w, magnetization transfer (MT) and multi-shell diffusion (DWI) MRI sequences. The detailed acquisition parameters for each image are summarized in Table 1 and data from two representative participants are represented in Figure S1.

**Table 1.** Acquisition parameters for each contrast image.

|  | T2*w | Magnetization<br>transfert (GRE-MT1 / GRE MT0) | DWI | T1w (MPRAGE) | EPI |
| --- | --- | --- | --- | --- | --- |
| **Structure coverage**<br>(top / bottom) | Brainstem / C7* | C1*-C7* | C1-C7* | Top of the brain /<br>T1* | Top of the brain /<br>T1* |
| **Coil elements** | HC: 1-7; NC: 1-2 | HC: 5-7; NC: 1-2 | HC: 7; NC: 1-2 | HC: 1-7; NC: 1-2 | HC: 1-7; NC: 1-2 |
| **Field of view** | 191x375 mm<sup>3</sup> | 241x168 mm<sup>3</sup> | 42x122 mm<sup>3</sup> | 364x375 mm<sup>3</sup> | 215x318 mm<sup>3</sup> |
| **Orientation** | Transversal | Transversal | Transversal | Transversal | Transversal |
| **Slices (n)** | 30 | 30 | 24 | 288 | 69 |
| **Volumes (n)** | - | - | - | - | 230 |
| **Voxel size** | 0.4x0.4x5 mm<sup>3</sup> | 0.9x0.9x5 mm<sup>3</sup> | 0.9x0.9x5 mm<sup>3</sup> | 1.3x1.3x1.3 mm<sup>3</sup> | 1.6x1.6x4 mm<sup>3</sup> |
| **iPAT mode** | Grappa | Grappa | - | Grappa | Grappa |
| **TR** | 34 ms | 35 ms | 780 ms | 2300 ms | 1550 ms |
| **TE** | 14 ms | 2.62 ms | 64 ms | 3.3 ms | 23 ms |
| **Flip angle** | 5 | 9 | 90 | 9 | 70 |
| **Acceleration factor**<br>(phase encoding) | 2 | 2 | - | 2 | 2 |
| **Multiband (slice**<br>**encoding)** | - | - | - | - | 3 |

### Data curation

Data were sorted, transformed into NIFTI, and organized using the Brain Imaging Data Structure (BIDS) standard (with dcm2bids v2.1.4). We used an in-house python pipeline (available here: https://github.com/CarolineLndl/Landelle_spinebrain_aging) based on the Spinal Cord Toolbox (SCT, version 5.6.0; De Leener et al., 2017), the Oxford Center for fMRI of the Software Library (FSL, version 5.0), the Statistical Parametric Mapping (SPM12, running on Matlab 2021b), the Tapas PhysiO toolbox (release 2022a, V8.1.0; Kasper et al., 2017) and the Nilearn toolbox (version 0.9.1).

### Structural data processing

After visual inspection of the structural images, three participants were excluded based upon T1w and T2*w analyses; five from the MTR and 12 for DWI for data quality issue. For analyses combining multiple contrasts, only the remaining common 54 participants were included (demographical details Table S1).

### Brain and spinal cord preprocessing

First, the T1w image was preprocessed, starting with the separation of brain and spinal cord slices to perform tailored preprocessing on each structure.

Brain tissue segmentation was performed automatically using CAT12 toolbox (an SMP12 extension) which employs the DARTEL (Diffeomorphic Anatomical Registration using Exponentiated Lie Algebra) algorithm to categorize brain voxels into gray matter (GW), white matter (WM) and cerebrospinal fluid (CSF) ^79^. A study-specific brain template in MNI space was generated from all participants’ brain T1w images using DARTEL. This study-specific template enables better inter-participant alignment by iteratively optimizing deformation fields to minimize anatomical variability, which is particularly beneficial when analyzing participants across a wide age range.

Spinal cord segmentation included the GM and WM, it was performed automatically (with SCT, *sct_propseg*), and manual adjustments were applied when necessary. First, the T1w image was warped into the PAM50 space (0.5 x 0.5 x 0.5 mm^3^) using cord segmentation and disk labelling (with SCT, *sct_register_to_template*). Next, spinal cord segmentation was computed on T2*w, MT0, MT1 (*i.e.*, with and without an MT pulse) and the mean DWI images by automatic methods (with SCT, sct_propseg) with manual corrections when necessary. Specific preprocessing was also applied to each contrast image. The GM of the T2*w images were automatically segmented (with SCT, sct_deepseg_gm), the MT0 image was registered to the MT1 image, and DWI images were corrected for motion correction (with SCT, sct_dwi_moco). Finally, the three images were normalized into the PAM50 template using their segmentation as well as the T1w to PAM50 warping field to initialize the registration (with SCT, *sct_register_multimodal*).

### Spinal cord microstructural metric extraction

We used the previously generated warping field to transform the PAM50 template to the individual space to extract metrics within spinal levels atlas (Frostell et al., 2016, SCT v6.1). Because of the limited reliability of automatic tissue segmentation at the C1 level—attributable to poor anatomical contrast—microstructural analyses were confined to segments C2 to C7 where tissue boundaries could be delineated more robustly. First, we calculated the number of voxels within the GM or the WM in the T2*w image (voxel counts) after automatic segmentation of the image in individual space. The magnetization transfer ratio (MTR) was computed for each voxel using the co-registered MT0 and MT1 images (*sct_compute_mtr)*. Finally, we computed fractional anisotropy (FA), mean diffusivity (MD), radial diffusivity (RD) and axial diffusivity (AD) from the DWI images using DIPY library *via* the SCT (*sct_dmri_compute_dti*). The average values of these microstructural metrics were extracted within the C2 to C7 spinal levels. Each spinal level was divided into four quadrants (ventral-left, ventral-right, dorsal-left, dorsal-right), resulting in a total of 24 spinal cord parcels (Fig. 1A). We choose to define levels based on spinal segment rather than vertebral levels to more accurately reflect the functional organization, which aligns with the rootlet insertion ^17^, thereby enabling a more robust comparison with our functional results.

### Spinal cord morphometric similarity matrix

We derived the spinal cord morphometric similarity matrix using the same procedure as that employed for the brain ^80^. Specifically, each of the 24 spinal regions was represented as a vector of multiple structural features (GM voxel counts, WM voxels counts, MTR, FA, MD, AD, RD) (Fig. 1b). Each of the feature vectors for each individual was normalized (z-scored). Pairwise Pearson correlations were computed as a measure of similarity between feature vectors of all parcel pairs to generate individual-level morphometric matrices (24×24). To estimate the within and between segmental networks similarity, the mean similarity was calculated for intra-segmental (*i.e*, the four parcels from the same spinal level) and inter-segmental (*i.e*, parcels from distinct spinal levels) parcel pairs. Finally, we average the matrices across individual to obtain a group-level morphometric matrix.

### Modeling age change in morphometric features

Given the strong structural asymmetric and distinct functional role between the ventral and dorsal divisions of the spinal cord (Landelle, et al., 2021), the age-related changes were estimated separately. Specifically, we estimated linear age-related changes in the average of each feature (not z-scored) across the ventral and dorsal parcels using an ordinary least squares (OLS) model that included age and sex as fixed effects. In addition, individual age values were predicted from these combined features in the ventral or dorsal parcels using an elastic net regression model (alpha = 1.2 and lambda = 1). To validate the model, we performed a 5-fold cross validation. In each iteration, the dataset was randomly split into 5 folds, with approximately 10-11 participants (20%) allocated for testing, while the remaining participants (80%) were used for training. This process was repeated 10 times, resulting in a total of 50 train-test evaluations. The mean absolute error (MAE) and the Pearson’s correlation coefficient (r) between predicted brain age and chronological age were averaged across the repetition to increase the robustness of the results. For each iteration, the model produces regression weights for each predictor, *i.e.* each feature, corresponding to the importance of each feature in predicting age.

### Brain and spinal cord structural metric extraction

To investigate the relationship between the structural integrity of the spinal cord and that of the brain for each individual, we computed spinal cord GM cross-sectional area (CSA) and brain GM volume. The motivation for using these two metrics is that they are the most widely used structural biomarkers for assessing spinal cord or brain atrophy, as they can be derived from various structural contrasts (T1w, T2w, MT) and are highly feasible in clinical settings. Here, we used the GM segmentation derived from T2*w images for the spinal cord to compute GM CSA and T1w images to calculate brain GM volume. Specifically, spinal cord structural integrity was assessed using GM cross-sectional area (CSA) measured for each of the six segmental levels from C2 to C7 (*via* SCT, *sct_process_segmentation*). By contrast, brain structural integrity was computed using total GM volume (in mL) across 200 GM cortical (Schaefer atlas 7Networks, Schaefer et al., 2018), 24 GM cerebellar and six subcortical structures (from the Cobra atlas, Tullo et al., 2018) extracted *via* the CAT12 toolbox. Finally, for each of the ten networks (seven Schaefer’s cortical, spinal cord, cerebellar and subcortical networks), we calculated the total structural integrity by summing brain GM volumes or by computing the mean GM-CSA across all corresponding parcels.

First, we estimated the relationship between spinal cord CSA and each brain network structure using an OLS model, with age and sex included as fixed effects. Then, we tested the interaction between brain network GM structure and age to determine whether age modulates this spinal cord-brain structure relationship. Age and sex fixed effects were also reported.

### Functional data processing

The 67 participants included for T1w and T2s analyses were also included for functional analyses (see Table S1 details about demographical population). Functional processes are similar to those developed in our previous studies ^17,18,20^.

### Preprocessing

Slice-timing correction was applied to the functional images, followed by the separation of brain and spinal cord slices to enable structure-specific preprocessing, which included the following steps:

*i) Motion correction*. Spinal cord motion correction used slice-wise realignment and spline interpolation (SCT, *sct_fmri_moco*) and was performed inside a 30 mm cylindrical mask centred on the spinal cord centerline,including the spinal canal. At the brain level, we removed the non-brain tissue FSL, BET) before performing motion correction with rigid-body realignment (FSL, MCFLIRT). Using the realignment motion parameters, we calculated the framewise displacement (*i.e., FD,* motion between two consecutive volumes) at both spinal cord and brain levels. Allparticipants had FD < 0.3 mm at the brain and spinal cord levels (Fig. S2B). Using OLS model using age as fixed effect, we found that the FD significantly increased with aging at the brain (t(65)= 4.75 p< 0.001) but not for the spinal cord (t(65)= 0.34 p=0.74, Fig. S2B). Then, we computed the temporal signal-to-noise ratio (tSNR) to evaluate the quality of the functional data over time. For each voxel, tSNR was calculated by dividing the mean signal intensity across time by its standard deviation. At the population level, we obtained a mean tSNR for within the spinal cord and within the brain (Fig. S2C). Using OLS model including age as fixed effect, we did not find significant age-related differences in the tSNR values for either the spinal cord (mean: 22.16 ± 2.0) ; t(65)= -1.70, p= 0.095) or brain (mean: 58.55 ± 4.9; t(65)= -1.24, p =0.22).
*ii) Image segmentation*. The resulting mean motion corrected image was segmented in two steps for the spinal cord: the spinal cord centerline was extracted manually, and the cord (GM + WM) and cerebrospinal fluid (CSF) were segmented (SCT, *sct_propseg*). Tissue segmentations were manually corrected when necessary.
*iii) Normalization into MNI or PAM50 template*. Similar to the microstructural images, the spinal cord mean motion corrected functional image was normalized into the PAM50 template using its cord segmentation as well as the T1w to PAM50 warping field to initialize the registration (with SCT, *sct_register_multimodal*). At the brain level, normalization was done in three steps using SPM12. First, we coregistered the mean functional image to the T1w space. Next, we warped the T1w image into the MNI template (2 x 2 x 2 mm^3^). Finally, the resulting deformation field was applied to move the functional images into MNI space. The functional to template warping fields were applied to the spinal cord and brain 4D functional images after the denoising steps.

### Time series denoising

We modelled slice-wise nuisance regressors to account for physiological noise for each participant. First, we used the RETROspective Image CORrection (RETROICOR) procedure ^81^ to computed noise regressors from peripheral physiological recordings (heart rate and respiration, Tapas PhysiO toolbox, an SPM extension, Kasper et al., 2017)). Specifically, we modeled four respiratory harmonics,three cardiac, and one multiplicative term for the interactions between respiratory and cardiac noise (18 regressors in total, similar to ^20,82^ and adjusted the regressors timing to each slice. Second, we used the CompCor approach ^83^ to identify non-neural fluctuations by extracting the first principal components of the unsmoothed brain or spinal cord cerebrospinal fluid (CSF) signal in the participant’s native space (12 components for the brain slices and 5 components for the spinal cord slices). The first five slice-wise discrete cosine transform (DCT) basis functions were added for detrending. These nuisance regressors were finally combined with the six brain and two spinal cord slice-wise motion parameters. The removal of the noise confounds was applied slice by slice and based on a projection on the orthogonal of the fMRI time-series space and was applied orthogonally to a band-pass temporal filter (0.01–0.17 Hz, with Nilearn, *img.clean_img*). No smoothing, nor any signal standardization was applied to the data at this step of the processing.

### Functional connectivity analyses

Functional connectivity was performed on the denoised time-series, normalized in template space and z-scored.

#### ICAPs framework

We employed the iCAPs framework to identify seven spinal cord segments in a data-driven manner ^17,84^. This approach extracts transient activity using total activation ^85^ and the temporal clustering of these signals. Specifically, a regularized hemodynamic deconvolution was used to extract activity-inducing signals from the smoothed, denoised, normalized, z-scored and unfiltered time series. Innovation signals (*i.e.,* temporal derivative of the activity-inducing time series) were then clustered using K-means to generate group-level innovation-driven coactivation patterns (iCAPs). To assess the spatial similarity with the Frostell’s atlas (Frostell et al., 2016), we computed the dice coefficient between the resulting components and the Frostell’s segmental levels.

#### Spinal cord functional connectivity analysis and age effects

The spinal cord was divided into four quadrants along the dorsal-ventral and left-right axes, across the seven spinal cord levels (*i.e.*, 28 parcels). For each individual, the unsmoothed and band-pass filtered time series within each parcel were extracted and averaged. The level of functional connectivity (FC) was then computed using Pearson’s correlation coefficient between each pair of parcel time-series, resulting in a 28×28 individual-level spinal functional connectivity (SpiFC) matrix. Individual matrices were then converted using Fisher’s r-to-z transformation. The population-level SpiFC was constructed as the mean SpiFC across all individuals. The mean FC was also calculated for intra-and inter-segmental level parcel pairs to assess intra-and inter-network FC. The age effect was assessed using an OLS regression model for intra-or inter-segmental levels of the six parcel pairs (ventro-ventral, dorso-dorsal, right dorso-ventral, left dorso-ventral, right dorso-left ventral and left dorso-right ventral), using age and sex as fixed effects. The statistical results were FDR-corrected for multiple comparisons.

#### Analysis of intra-vs. inter FC and age effects across the brain and spinal cord

To assess whether the spinal cord FC organization is similar to that of the brain, we conducted parallel analyses at the brain level using data from the same individuals. The brain was parcelled into 200 cortical areas using the seven-network Schaefer atlas ^45^, along with 24 cerebellar and six subcortical regions using the Cobra atlas ^46^, yielding a 230×230 individual-level brain functional connectivity (BrainFC) matrix. The brain parcels were grouped into cerebellar, subcortical and 7 cortical networks. To evaluate the FC organization across the CNS, we computed intra-and inter-network FC separately within the spinal cord (averaged across the seven spinal segments, each treated as a distinct network) and the brain (one subcortical, one cerebellar and seven cortical networks). This parallel approach allowed us to investigate systematically inter-*vs.* intra-network FC across CNS regions and examine whether the spinal cord follows similar network organization principles as the brain does. Finally, we investigated the age-related changes in this FC organization. Specifically, intra-and inter-network FC for each the brain and spinal cord network were entered in an OLS regression model that included age and sex as fixed effects. The statistical results were FDR-corrected for multiple comparisons.

### Time-series dynamic properties extraction

Time-series dynamic properties extraction was performed on time-series denoised, unsmoothed and normalized in template space.

#### CAnonical Time-series Characteristics (catch-22) extraction

For a given spinal cord parcel (28 parcels, from C1 to C7 segmental levels divided in four quadrants), the dynamic properties of the z-scored BOLD time-series were summarized using a set of features extracted with the Canonical Time-series Characteristics python toolbox, catch22 ^44^. The catch-22 features were based on the hctsa toolbox ^86^ and designed to be the highly explanatory subset of features typically encountered in time-series (while not specifically defined for BOLD signal). We excluded five features that exhibited a dominant single value in more than 25% of the observations (*i.e.,* across spinal parcels and participants), as such features inherently lack variability and discriminative, violated the assumptions that features capture meaningful variability (Fig. S7). We also add to these features the calculation of the standard deviation and the mean on no-zscored time-series. To compare our data with a standard biomarker commonly used at the spinal cord level ^41–43^ to assess local resting-state activity, we calculated the amplitude of low-frequency fluctuations (ALFF). ALFF was computed by first deriving the power spectrum of the detrended non-zscored BOLD time series at each voxel, then taking the square root of the mean power within the low-frequency range (0.01– 0.027 Hz). The average ALFF value was then calculated within each of the 28 spinal cord parcels. In total, we obtained 20 time-series dynamic features (17 from catch-22, mean, standard deviation, ALFF) that include, but are not limited to, distributional properties, autocorrelations and variability of a given time-series.

#### Spinal cord dynamic profile similarity matrix (SpiDyn)

To compute the dynamic profile similarity matrix across the 28 spinal parcels, each feature was normalized across spinal cord parcels to the unit interval using a scaled robust sigmoid function, ensuring comparability across features with different value ranges ^87^. Similar to the approach used for computing the morphometric matrices, the spinal cord dynamic profile similarity matrices (SpiDyn matrices) were generated by calculating pairwise Pearson correlations between the 19-dimensional feature vectors across all parcel pairs, resulting in individual-level 28×28 similarity matrices (see details in *Spinal cord morphometric similarity matrix* section).

#### Modelling age change in time-series dynamic features

When analyzing each time-series dynamic feature to test the age effect, normalization or rescaling was not applied to preserve interpretability in the original feature scale and avoid introducing distortions. We estimated linear age-related changes in the average of each feature across right-ventral, left-ventral, right-dorsal and left-dorsal parcels using an OLS model including age and sex as fixed effects. The statistical results were FDR-corrected for multiple comparisons.

#### Analysis of local time-series dynamics and age effects across the brain and spinal cord

To assess the topographical organization of time-series dynamics across the CNS, we applied principal component analysis (PCA; n=10 components) to the parcels-by-features matrix (254 parcels x 20 features) to capture independent patterns of time-series dynamics. The resulting component loadings were summarized by computing the centroid for each of the predefined networks (nine brain and two spinal cord networks). For this analysis, the spinal cord was subdivided into ventral and dorsal networks (encompassing, respectively, left and right ventral or dorsal parcels) to test whether an intrinsic functional dissociation or association exists between these subdivisions, analogous to the antero/posterior or unimodal/transmodal gradients observed at the cortical level ^5,49^. Through this analysis, we aimed to uncover the gradient of functional dynamics spanning the whole CNS (cortical, subcortical and spinal cord), thereby revealing the large-scale organization principle.

To assess age-related changes in local dynamics, we average each parcel’s feature within their assigned networks (nine brain and two spinal cord networks) and performed OLS regression models, for each network and feature, using age and sex as fixed effects. The statistical results were FDR-corrected for multiple comparisons.

### Structural and functional coupling

We estimated regional structure-SpiFC, structure-SpiDyn and SpiFC-SpiDyn coupling by computing Spearman correlation between each parcel’s profile in the respective matrices (i.e, correlations between rows for each region). Then we investigated the age-related change in these couplings. Specifically, we applied OLS regression models for each spinal region, with coupling strength as the dependent variable, age as a fixed effect and sex as a covariate.

Finally, to determine whether spinal regions exhibiting age-related changes in structural, SpiFC or SpiDyn also show the corresponding effects in their coupling, we computed Spearman’s correlations between t-values from the OLS models across regions.

## Data and Code Availability

The datasets generated and analyzed during the current study are available upon request. The code used for data pre-processing and analysis is available for public use here: https://github.com/CarolineLndl/Landelle_spinebrain_aging.

## Author contributions

C.L. and J.D. conceptualized the study. C.L. collected the data, performed the preprocessing and formal analysis. C.L., B.DL., N.K., and S.S. developed the methodology. C.L. created the figures and drafted the manuscript. All authors reviewed and edited the manuscript J.D. supervised the project and secured the funding.

## Acknowledgments

This work was performed at the McConnell Brain Imaging Centre, The Neuro (McGill University, Montreal, Canada). We thank David Costa, Ronaldo Lopez and Soheil Mollamohseni Quichani for their help in data acquisition and all participants who were recruited for this study. C.L was supported by the Canadian Institutes of Health Research (CIHR, Grant. No. 13556932). J.D. has received grants to support the work from “Fondation Courtois”, the Natural Sciences and Engineering Research Council of Canada (NSERC, Grant No. 248074); Brain Canada Platform (Grant No. 255934) and Healthy Brain for Healthy Lives (HBHL, Grant No. 2473633). J.D, B.DL and V.MP received funding from the STRATALS EU Joint Programme – Neurodegenerative Disease Research (JPND) 2019. N.K. and D.V.D.V. were supported by the Swiss National Science Foundation under Project No. 205321_207493. The funders had no role in study design, data collection and analysis, decision to publish, or preparation of the manuscript.

