## Supplementary material for "Spinal cord structural and functional architecture and its shared organization with the brain across the adult lifespan"

### Supplementary figures

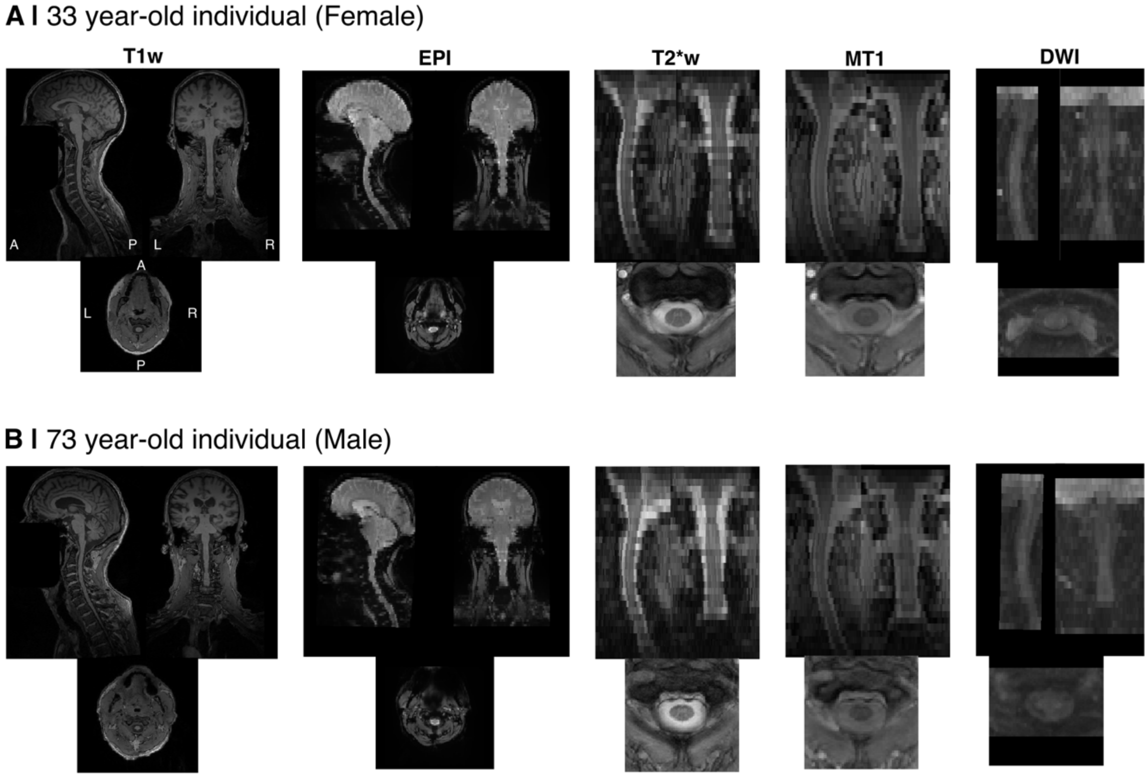


**Fig. S1 | Representative individuals**. Example of raw images for T1-weighted (T1w), T2*-weighted (T2*w), Magnetization transfer (MT1), and mean images for EPI, Diffusion weighted images (DWI) of two participants: (A) 33 year-old female, (B) 73 year-old male)

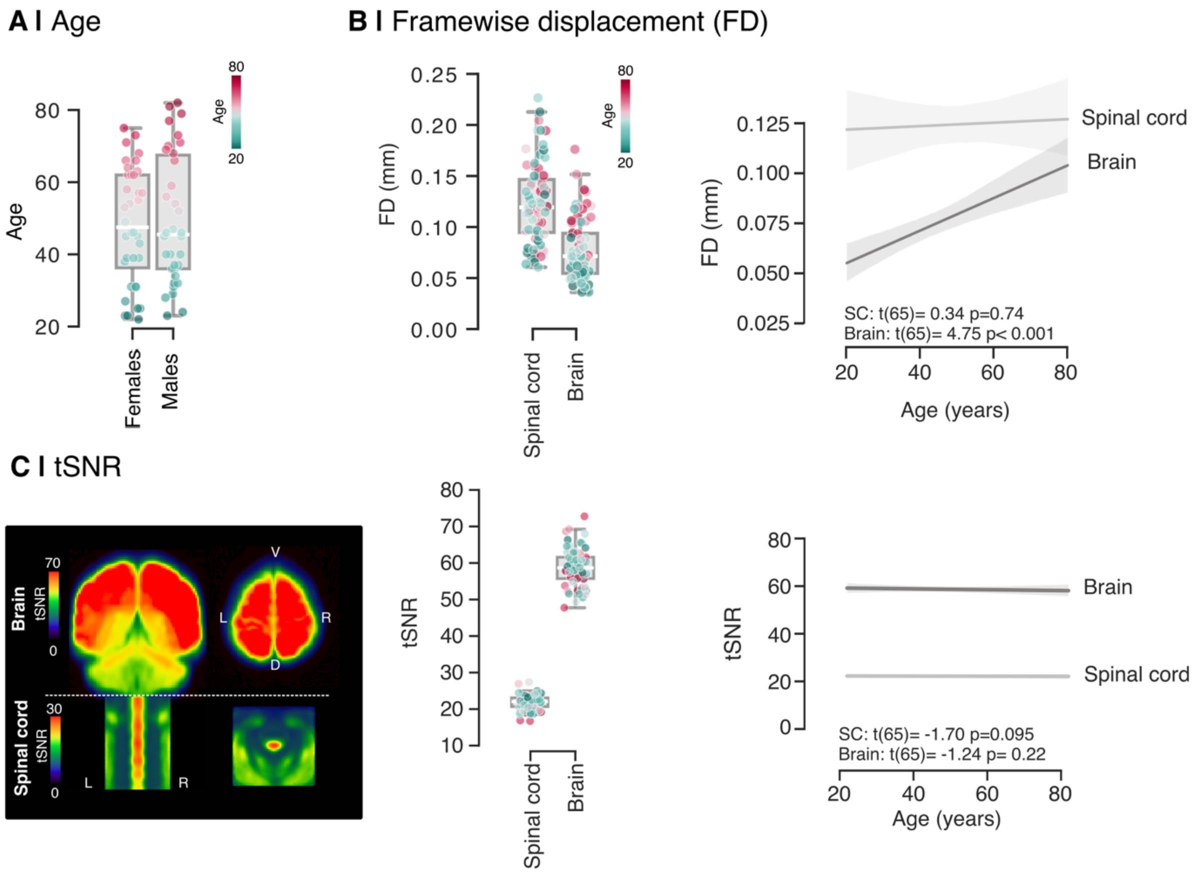


**Figure S2 | Age distribution and quality check metrics. A.** Distribution of participant ages by sex. **B.** Framewise displacement (FD). Left: boxplots of FD for spinal cord and brain masks. Each box extends from the 25th to the 75th percentile of the group's distribution and the medians are represented by the horizontal line inside the box. The vertical extending lines denote the extreme values within 1.5 interquartile range. Individual data points are colored by participant age from green (younger) to pink (older). Right: Linear regressions between age and FD for the spinal cord and the brain (right) **C.** Temporal signal-to-noise ratio (tSNR). Left: Average tSNR maps for the brain and the spinal cord, display with distinct color scales for optimal visualization. Middle: boxplots of tSNR for the spinal cord and brain masks. Each box extends from the 25th to the 75th percentile of the group's distribution and the medians are represented by the horizontal line inside the box. Individual data points are colored by participant age from green (younger) to pink (older). Right: Linear regressions between age and tSNE for the spinal cord and the brain
D: dorsal; V: ventral; L: left; R: right.

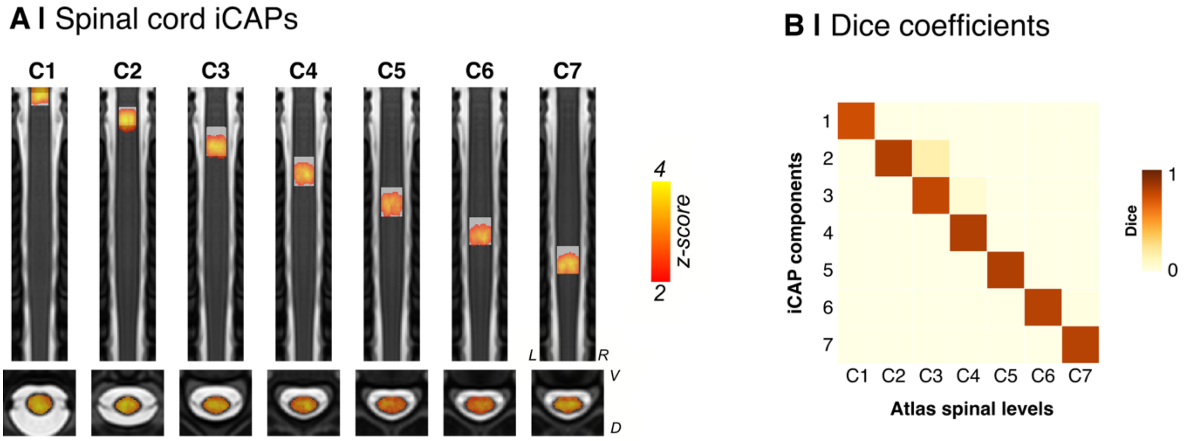


**Figure S3 | Spinal cord segmental levels. A.** Spinal cord iCAPs presented in rostro-caudal order. The label of the iCAPs (from C1 to C7) are determined based on the spinal segmental atlas (Frostell et al., 2016), plotted as a white underlay. L: left; R: right; V: ventral; D: dorsal. **B.** The similarity between the iCAP components and the atlas-based spinal levels was evaluated using Dice coefficients between the two maps.

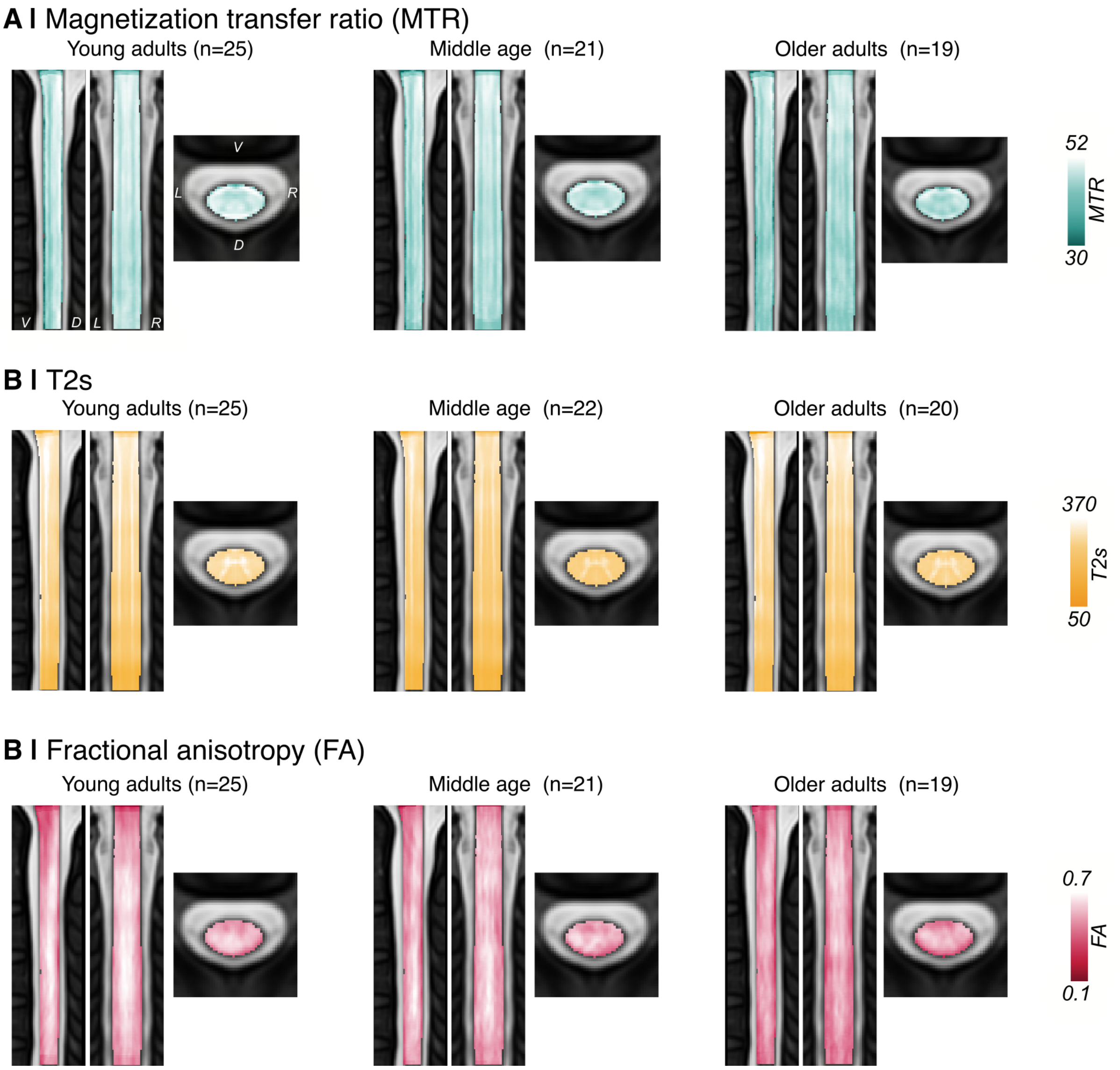


**Figure S4 | Average metric maps.** We computed the average metric maps for MTR (A), T2s (B) and FA (C) metrics by dividing into three age groups: young adults (20-40 years), middle-aged adults (40-60 years) and older adults (60-80 years). L: left; R: right; V: ventral; D: dorsal, MTR: Magnetization transfer ratio, FA: Fractional anisotropy

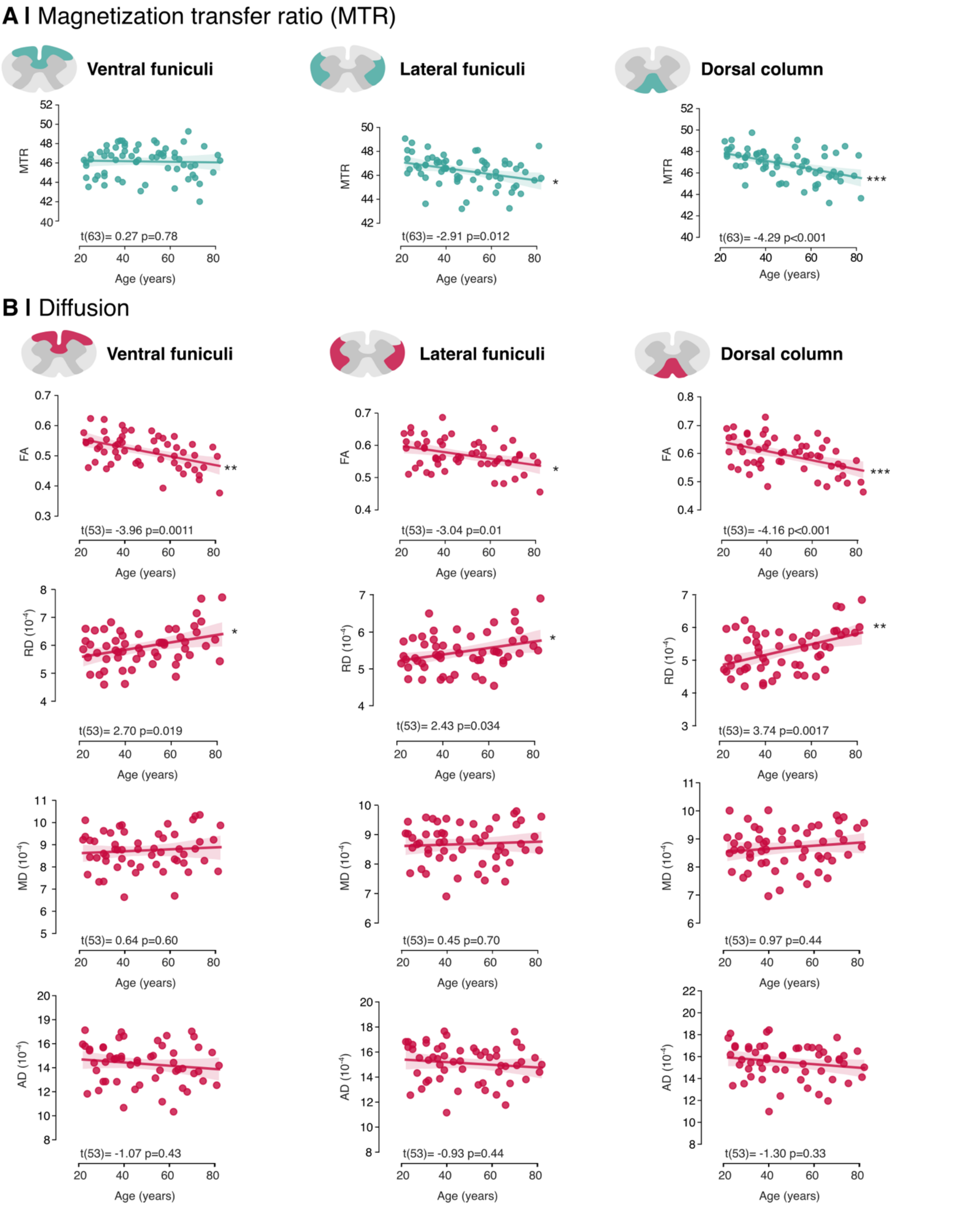


**Fig S5 | Microstructural changes in the spinal cord white matter (WM) tracts.** Magnetization transfer ratio (MTR, **A**) and several diffusion metrics (**B**) were extracted within three white matter tracts (dorsal column, lateral funiculi and ventral funiculi). Linear regressions between age and each metric were computed using ordinary least squares (OLS), with age and sex as fixed factors. t-values and degree of freedom (t(df)) as well as FDR-corrected p-values for age effect, are reported on each graph. FA: fractional anisotropy, RD: radial diffusivity, MD: mean diffusivity, AD: axial diffusivity. P-values are corrected for multiple comparison *: $p < 0.05$; ***: $p < 0.01$; ***: $p < 0.001$;

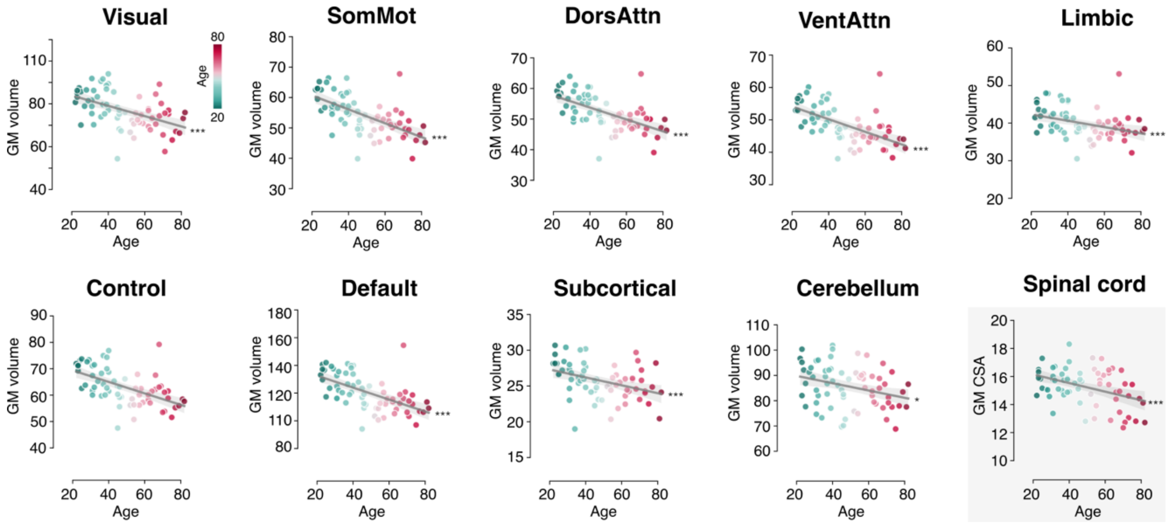


**Fig S6 | Structural changes in the gray matter (GM).** GM volume was extracted across seven cortical networks, as well as the cerebellum and subcortical regions. For the spinal cord, cross-sectional area (CSA) was measured. Linear regressions between age and each metric were computed using ordinary least squares (OLS), with age and sex as fixed factors. Age effect: *: $p_{FDR} < 0.05$; ***: $p_{FDR} < 0.01$; ***: $p_{FDR} < 0.001$.

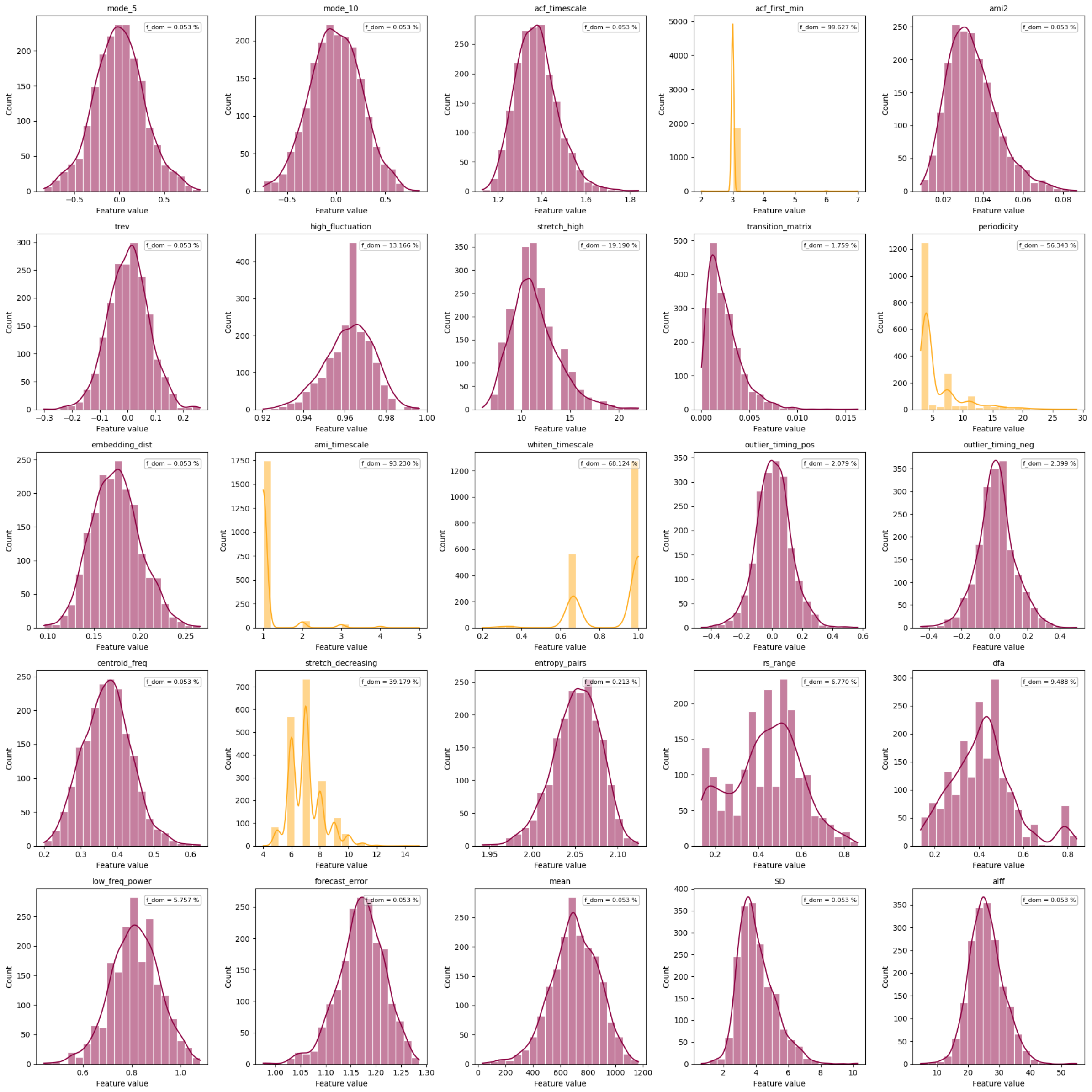


**Figure S7 | Distribution of the SpiDyn features.** Distribution of each feature, with pink indicating the features retained for further analyses and yellow indicating features excluded due to a dominant value and inherently lack of variability. The proportion of dominant value is reported on each graph as ‘f_dom’.

### Supplementary tables

| Contrasts | # Participants (#Females/ #Males) | Age |
|---|---|---|
| fMRI/ T1w /T2s | 67 (36/31) | 48.67 ± 17.18 |
| MTR | 65 (35/30) | 48.28 ± 17.08 |
| DWI | 55 (29/26) | 48.29 ± 17.58 |
| All | 54 (28/26) | 48.33 ± 17.74 |

**Table S1 | Demographic population.** The number of participants, mean ± standard deviation as well as the number of females and males, are provided for each contrast after participant exclusions. The last row evaluates the population across the different contrasts

| Contrast | ROI | Fixed effect | t-value | p-value | p-value FDR | Significance |
|---|---|---|---|---|---|---|
| **MTR (n=55)** | **DL** | age | -3.88 | <0.001 | <0.001 | *** |
| | | sex | -0.008 | 0.99 | <0.99 | ns |
| | **DR** | age | -3.43 | <0.001 | 0.0017 | ** |
| | | sex | 0.11 | 0.90 | 0.95 | ns |
| | **VL** | age | -1.22 | 0.22 | 0.30 | ns |
| | | sex | -2.53 | 0.012 | 0.030 | * |
| | **VR** | age | -0.31 | 0.75 | 0.82 | ns |
| | | sex | -0.98 | 0.32 | 0.48 | ns |
| **GM (n=67)** | **DL** | age | -4.05 | <0.001 | <0.001 | *** |
| | | sex | 3.87 | <0.001 | <0.001 | *** |
| | **DR** | age | -3.63 | <0.001 | <0.001 | *** |
| | | sex | 3.82 | <0.001 | <0.001 | *** |
| | **VL** | age | -2.63 | 0.009 | 0.017 | * |
| | | sex | 3.62 | <0.001 | 0.015 | * |
| | **VR** | age | -2.23 | 0.026 | 0.045 | * |
| | | sex | 3.43 | <0.001 | 0.002 | ** |
| **WM (n=67)** | **DL** | age | -0.10 | 0.91 | 0.91 | ns |
| | | sex | 0.10 | 0.92 | 0.95 | ns |
| | **DR** | age | -1.37 | 0.17 | 0.26 | ns |
| | | sex | -0.18 | 0.86 | 0.95 | ns |
| | **VL** | age | -0.62 | 0.53 | 0.59 | ns |
| | | sex | -0.47 | 0.63 | 0.84 | ns |
| | **VR** | age | -0.17 | 0.90 | 0.91 | ns |
| | | sex | -0.32 | 0.75 | 0.91 | ns |
| **FA** | **DL** | age | -6.34 | <0.001 | <0.001 | *** |

| | | | | | | |
|---|---|---|---|---|---|---|
| (n=55) | | sex | -4.21 | <0.001 | <0.001 | *** |
| | DR | age | -6.00 | <0.001 | <0.001 | *** |
| | | sex | -5.50 | <0.001 | <0.001 | *** |
| | VL | age | -5.08 | <0.001 | <0.001 | *** |
| | | sex | -1.88 | 0.061 | 0.11 | ns |
| | VR | age | -4.40 | <0.001 | <0.001 | *** |
| | | sex | 2.50 | 0.013 | 0.03 | * |
| **RD (n=55)** | **DL** | age | 4.73 | <0.001 | <0.001 | *** |
| | | sex | 2.90 | <0.001 | 0.0040 | * |
| | **DR** | age | 4.20 | <0.001 | <0.001 | *** |
| | | sex | 4.50 | <0.001 | <0.001 | *** |
| | **VL** | age | 3.70 | <0.001 | <0.001 | *** |
| | | sex | -0.36 | 0.061 | 0.71 | ns |
| | **VR** | age | 2.95 | 0.0034 | 0.0080 | ** |
| | | sex | 0.58 | 0.56 | 0.78 | ns |
| **AD (n=55)** | **DL** | age | -2.64 | 0.0087 | 0.017 | * |
| | | sex | -2.41 | 0.017 | 0.036 | * |
| | **DR** | age | -2.45 | 0.015 | 0.027 | * |
| | | sex | -2.08 | 0.039 | 0.072 | . |
| | **VL** | age | -1.20 | 0.23 | 0.31 | ns |
| | | sex | -3.25 | 0.0013 | 0.0044 | ** |
| | **VR** | age | -1.44 | 0.15 | 0.25 | ns |
| | | sex | -2.83 | 0.0054 | 0.014 | * |
| **MD (n=55)** | **DL** | age | 1.08 | 0.28 | 0.36 | ns |
| | | sex | 0.15 | 0.87 | 0.95 | ns |
| | **DR** | age | 0.75 | 0.45 | 0.54 | ns |
| | | sex | 1.18 | 0.24 | 0.37 | ns |
| | **VL** | age | 1.24 | 0.21 | 0.31 | ns |
| | | sex | -2.22 | 0.027 | 0.054 | . |
| | **VR** | age | 0.73 | 0.46 | 0.54 | ns |
| | | sex | -1.42 | 0.16 | 0.26 | ns |

**Table S2 | Age and sex effects on individual morphometric features.** The age-related effect was obtained using an ordinary least squares (OLS) model including age and sex as fixed effects. The fixed effects of each model (each feature and each quadrant) is reported. The number of participants included in each model is reported in the first column (n), and the t, p-values and FDR-corrected p-values are reported for each fixed effect. For the sex effect, Females is the reference level. The coefficient (and its t-value) represents the extent to which the mean outcome for Males differs from that for Females; thus, a positive value indicates higher values in Males *compared to* Females, and *vice versa*. The last column represents the significance: ns: not significant, . : significant before FDR but not after, *: $p_{FDR} < 0.05$, **: $p_{FDR} < 0.01$; ***: $p_{FDR} < 0.001$.

| ROI-pairs | Fixed effect | t-value | p-value | p-value FDR | Significance |
|---|---|---|---|---|---|
| **Dorso-Dorsal (DL-DR)** | age | 1.62 | 0.10 | 0.10 | ns |
| | sex | -2.71 | 0.0069 | 0.040 | * |
| **Ventro-Ventral (VL-VR)** | age | 2.43 | 0.014 | 0.018 | * |
| | sex | -1.24 | 0.21 | 0.32 | ns |
| **R Dorso-Ventral (DR-VR)** | age | 2.71 | 0.0068 | 0.014 | * |
| | sex | -0.86 | 0.38 | 0.40 | ns |
| **L Dorso-Ventral (DL-VL)** | age | 2.55 | 0.010 | 0.016 | * |
| | sex | -0.83 | 0.40 | 0.40 | ns |
| **L Cross (DL-VR)** | age | 3.62 | <0.001 | 0.0017 | ** |
| | sex | -1.26 | 0.21 | 0.32 | ns |
| **R Cross (DR-VL)** | age | 2.94 | 0.003 | 0.0096 | ** |
| | sex | -1.90 | 0.056 | 0.17 | ns |

**Table S3 | Age and sex effects on inter-segmental functional connectivity.** The age-related effect was obtained using an ordinary least squares (OLS) model including age and sex as fixed effects. The fixed effects of each model (each pair of ROIs between distinct segments) is reported. The degree of freedom was always 65 (i.e., 67 individuals – 2 fixed factors) and the t, p-values and FDR-corrected p-values are reported for each fixed effect. For the sex effect, Females is the reference level. The coefficient (and its t-value) represents the extent to which the mean outcome for Males differs from that for Females; thus, a positive value indicates higher values in Males *compared to* Females, and *vice versa*. The last column represents the significance: ns: not significant, . : significant before FDR but not after, *: $p_{FDR} < 0.05$, **: $p_{FDR} < 0.01$; ***: $p_{FDR} < 0.001$. DL: dorsal left, DR: dorsal right, VL: ventral left, VR: ventral right

| ROI-pairs | Fixed effect | t-value | p-value | p-value FDR | Significance |
|---|---|---|---|---|---|
| **Dorso-Dorsal (DL-DR)** | age | -0.10 | 0.92 | 0.92 | ns |
| | sex | -2.048 | 0.040 | 0.048 | * |
| **Ventro-Ventral (VL-VR)** | age | -1.80 | 0.072 | 0.14 | ns |
| | sex | -3.89 | <0.001 | <0.001 | *** |
| **R Dorso-Ventral (DR-VR)** | age | 1.57 | 0.11 | 0.17 | ns |
| | sex | -1.79 | 0.072 | 0.072 | ns |
| **L Dorso-Ventral (DL-VL)** | age | 2.71 | 0.0065 | 0.039 | * |
| | sex | -2.36 | 0.018 | 0.027 | * |
| **L Cross (DL-VR)** | age | 2.26 | 0.023 | 0.069 | . |
| | sex | -2.72 | 0.0064 | 0.019 | * |
| **R Cross (DR-VL)** | age | 1.44 | 0.15 | 0.18 | ns |
| | sex | -2.39 | 0.017 | 0.072 | . |

**Table S4 | Age and sex effects on intra-segmental functional connectivity.** The age-related effect was obtained using an ordinary least squares (OLS) model including age and sex as fixed effects. The fixed effects of each model (each pair of rois within the same segment) is reported. The degree of freedom was always 65 (i.e., 67 individuals – 2 fixed factors) and the t, p-values and FDR-corrected p-values are reported for each fixed effect. For the sex effect, Females is the reference level. The coefficient (and its t-value) represents the extent to which the mean outcome for Males differs from that for Females; thus, a positive value indicates higher values in Males *compared to* Females, and *vice versa*. The last column represents the significance: ns: not significant, . : significant before FDR but not after, *: $p_{FDR} < 0.05$, **: $p_{FDR} < 0.01$; ***: $p_{FDR} < 0.001$. DL: dorsal left, DR: dorsal right, VL: ventral left, VR: ventral right

| Feature | Rois | Fixed effect | t-value | p-value | p-value FDR | Significance |
|---|---|---|---|---|---|---|
| SD | DL | age | 3.71 | <0.001 | 0.016 | * |
| | | sex | 1.44 | 0.15 | 0.74 | ns |
| | DR | age | 3.20 | <0.001 | 0.035 | * |
| | | sex | 1.72 | 0.85 | 0.64 | ns |
| | VL | age | 2.11 | 0.035 | 0.24 | . |
| | | sex | 0.30 | 0.76 | 0.99 | ns |
| | VR | age | 0.80 | 0.42 | 0.73 | ns |
| | | sex | 0.033 | 0.97 | 0.99 | ns |
| ACF timescale | DL | age | 3.12 | 0.0018 | 0.036 | * |
| | | sex | -0.84 | 0.40 | 0.89 | ns |
| | DR | age | 1.72 | 0.085 | 0.42 | ns |
| | | sex | 0.93 | 0.35 | 0.89 | ns |
| | VL | age | 1.65 | 0.099 | 0.42 | ns |
| | | sex | 0.077 | 0.93 | 0.99 | ns |
| | VR | age | 1.60 | 0.11 | 0.42 | ns |
| | | sex | -1.60 | 0.11 | 0.63 | ns |
| ALFF | DL | age | 2.90 | 0.0037 | 0.042 | * |
| | | sex | 4.47 | <0.001 | <0.001 | *** |
| | DR | age | 2.70 | 0.0070 | 0.071 | . |
| | | sex | 4.04 | <0.001 | 0.002 | ** |
| | VL | age | 1.036 | 0.29 | 0.66 | ns |
| | | sex | 3.02 | 0.0025 | 0.068 | . |
| | VR | age | 0.58 | 0.56 | 0.77 | ns |
| | | sex | 2.89 | 0.0038 | 0.077 | . |
| Centroid frequency | DL | age | 3.34 | <0.001 | 0.034 | * |
| | | sex | -0.34 | 0.73 | 0.99 | ns |
| | DR | age | 1.01 | 0.31 | 0.65 | ns |
| | | sex | 1.15 | 0.25 | 0.82 | ns |
| | VL | age | 1.49 | 0.13 | 0.43 | ns |
| | | sex | -0.025 | 0.97 | 0.99 | ns |
| | VR | age | 1.57 | 0.11 | 0.42 | ns |
| | | sex | -1.55 | 0.12 | 0.64 | ns |
| Embedding distance | DL | age | 2.07 | 0.03 | 0.24 | . |
| | | sex | -0.69 | 0.49 | 0.95 | ns |
| | DR | age | 1.37 | 0.16 | 0.48 | ns |
| | | sex | 0.48 | 0.63 | 0.87 | ns |
| | VL | age | 1.07 | 0.71 | 0.66 | ns |
| | | sex | -0.37 | 0.71 | 0.99 | ns |
| | VR | age | 2.01 | 0.04 | 0.25 | . |
| | | sex | -2.64 | 0.008 | 0.13 | . |
| | DL | age | -2.21 | 0.027 | 0.21 | . |

| | | | | | | |
|---|---|---|---|---|---|---|
| | | sex | 0.85 | 0.39 | 0.89 | ns |
| Entropy pairs | DR | age | -0.75 | 0.45 | 0.72 | ns |
| | | sex | -1.26 | 0.21 | 0.82 | ns |
| | VL | age | -0.82 | 0.41 | 0.73 | ns |
| | | sex | -0.12 | 0.90 | 0.99 | ns |
| | VR | age | -0.077 | 0.94 | 0.95 | ns |
| | | sex | -1.77 | 0.077 | 0.64 | ns |
| Low frequency power | DL | age | -2.94 | 0.0032 | 0.042 | * |
| | | sex | 0.28 | 0.77 | 0.99 | ns |
| | DR | age | -1.68 | 0.09 | 0.42 | ns |
| | | sex | -1.63 | 0.10 | 0.63 | ns |
| | VL | age | -1.28 | 0.20 | 0.53 | ns |
| | | sex | -0.55 | 0.58 | 0.97 | ns |
| | VR | age | -1.58 | 0.11 | 0.42 | ns |
| | | sex | 0.86 | 0.38 | 0.89 | ns |
| Forecast error | DL | age | -3.058 | 0.0022 | 0.035 | * |
| | | sex | 0.35 | 0.72 | 0.99 | ns |
| | DR | age | -2.066 | 0.039 | 0.23 | . |
| | | sex | -0.50 | 0.61 | 0.97 | ns |
| | VL | age | -1.75 | 0.079 | 0.42 | ns |
| | | sex | -0.053 | 0.96 | 0.99 | ns |
| | VR | age | -3.38 | 0.017 | 0.15 | . |
| | | sex | 1.10 | 0.27 | 0.82 | ns |

**Table S5 | Age and sex effects on functional dynamic profiles.** The age-related effect was obtained for each feature and four regions using an ordinary least squares (OLS) model including age and sex as fixed effects. We report the fixed effects of the model when the age effect was significant before FDR correction for at least one region. The degree of freedom was always 65 (*i.e.,* 67 individuals – 2 fixed factors) and the t, p-values and FDR-corrected p-values are reported for each fixed effect. For the sex effect, Female is the reference level. The coefficient (and its t-value) represents the extent to which the mean outcome for Males differs from that for Females; thus, a positive value indicates higher values in Males *compared to* Females, and *vice versa*. The last column represents the significance: ns: not siginifcant, . : significant before FDR but not after, *: $p_{FDR} < 0.05$, **: $p_{FDR} < 0.01$; ***: $p_{FDR} < 0.001$. DL: dorsal left, DR: dorsal right, VL: ventral left, VR: ventral right, SD: Standard deviation, ALFF: amplitude of low frequency fluctuation, ACF: autocorrelation function.

| Networks | Fixed effect | t-value | p-value | p-value FDR | Significance |
|---|---|---|---|---|---|
| **Visual** | age | -4.73 | <0.001 | <0.001 | *** |
| | sex | 2.02 | 0.047 | 0.089 | . |
| **SomMot** | age | -6.49 | <0.001 | <0.001 | *** |
| | sex | 1.97 | 0.052 | 0.089 | ns |
| **DorsAttn** | age | -6.05 | <0.001 | <0.001 | *** |
| | sex | 1.88 | 0.064 | 0.092 | ns |
| **VentAttn** | age | -6.11 | <0.001 | <0.001 | *** |
| | sex | 2.36 | 0.021 | 0.070 | . |
| **Limbic** | age | -3.66 | <0.001 | <0.001 | *** |
| | sex | 4.13 | <0.001 | 0.0011 | ** |
| **Control** | age | -5.54 | <0.001 | <0.001 | *** |
| | sex | 2.17 | 0.033 | 0.084 | . |
| **Default** | age | -6.10 | <0.001 | <0.001 | *** |
| | sex | 2.69 | 0.009 | 0.046 | * |
| **Subcortical** | age | -3.50 | <0.001 | <0.001 | *** |
| | sex | 1.75 | 0.084 | 0.10 | ns |
| **Cerebellum** | age | -2.62 | 0.011 | 0.011 | * |
| | sex | 1.33 | 0.19 | 0.21 | ns |
| **Spinal cord** | age | -3.52 | <0.001 | <0.001 | *** |
| | sex | -0.90 | 0.37 | 0.37 | ns |

**Table S6 | Age and sex effects on brain GM volume and spinal cord CSA.** The age-related effect on gray matter (GM) volume was obtained for each brain network and the GM cross-sectional areas for the spinal cord using an ordinary least squares (OLS) model including age and sex as fixed effects. The degree of freedom was always 65 (*i.e.,* 67 individuals – 2 fixed factors) and the t, p-values and FDR-corrected p-values are reported for each fixed effect. For the sex effect, Female is the reference level. The coefficient (and its t-value) represents the extent to which the mean outcome for Males differs from that for Females; thus, a positive value indicates higher values in Males *compared to* Females, and *vice versa*. The last column represents the significance: ns: not significant, . : significant before FDR but not after, *: $p_{FDR} < 0.05$, **: $p_{FDR} < 0.01$; ***: $p_{FDR} < 0.001$.

| Networks | Intra FC (Mean ± SD) | Inter (Mean ± SD) | t-value | p-value | Significance |
|---|---|---|---|---|---|
| **Visual** | 0.20 ± 0.074 | 0.0086 ± 0.015 | 13.20 | <0.001 | *** |
| **SomMot** | 0.16 ± 0.051 | 0.022 ± 0.017 | 23.09 | <0.001 | *** |
| **DorsAttn** | 0.092 ± 0.034 | 0.020 ± 0.019 | 18.60 | <0.001 | *** |

| | | | | | |
|---|---|---|---|---|---|
| **VentAttn** | 0.13 ± 0.044 | 0.031 ± 0.015 | 20.78 | <0.001 | *** |
| **Limbic** | 0.13 ± 0.044 | 0.025 ± 0.016 | 21.31 | <0.001 | *** |
| **Control** | 0.067 ± 0.028 | 0.016 ± 0.015 | 14.95 | <0.001 | *** |
| **Default** | 0.095 ± 0.040 | 0.013 ± 0.015 | 16.62 | <0.001 | *** |
| **Subcortical** | 0.27 ± 0.081 | 0.029 ± 0.015 | 24.67 | <0.001 | *** |
| **Cerebellum** | 0.16 ± 0.069 | 0.030 ± 0.018 | 18.30 | <0.001 | *** |
| **Spinal cord** | 0.17 ± 0.077 | 0.033 ± 0.024 | 17.25 | <0.001 | *** |

**Table S7 | Intra *versus* inter-networks functional connectivity.** The comparison between intra *versus* inter-network functional connectivity was obtained using a paired–t-test. The degree of freedom was always 66 and the mean ± standard deviation, t-values and p-values are reported. The last column represents the significance: ns: not significant, *: p< 0.05, **: p< 0.01; ***: p < 0.001. SD: standard deviation

| Networks | Intra FC | | | Inter FC | | |
|---|---|---|---|---|---|---|
| | t-value | p-value | Signif. | t-value | p-value | Signif. |
| **Visual** | 13.10 | <0.001 | *** | 4.85 | <0.001 | *** |
| **SomMot** | 25.11 | <0.001 | *** | 10.56 | <0.001 | *** |
| **DorsAttn** | 20.43 | <0.001 | *** | 9.015 | <0.001 | *** |
| **VentAttn** | 24.50 | <0.001 | *** | 16.53 | <0.001 | *** |
| **Limbic** | 23.33 | <0.001 | *** | 12.27 | <0.001 | *** |
| **Control** | 19.40 | <0.001 | *** | 8.33 | <0.001 | *** |
| **Default** | 19.70 | <0.001 | *** | 7.00 | <0.001 | *** |
| **Subcortical** | 26.90 | <0.001 | *** | 15.37 | <0.001 | *** |
| **Cerebellum** | 19.04 | <0.001 | *** | 13.54 | <0.001 | *** |
| **Spinal cord** | 18.37 | <0.001 | *** | 11.40 | <0.001 | *** |

**Table S8 | Intra and inter-networks functional connectivity.** The comparison between functional connectivity and 0 was obtained using a t-test against 0. The degree of freedom was always 66 and the t-values and p-values are reported. The last column represents the significance: ns: not significant, *: p< 0.05, **: p< 0.01; ***: p < 0.001. SD: standard deviation

| Networks | Fixed effect | t-value | p-value | p-value FDR | Significance |
|---|---|---|---|---|---|
| **Visual** | age | 3.19 | 0.0022 | 0.0055 | ** |
| | sex | -2.98 | 0.0040 | 0.016 | * |
| **SomMot** | age | 2.50 | 0.015 | 0.016 | * |
| | sex | -2.43 | 0.018 | 0.035 | * |

| | | | | | |
|---|---|---|---|---|---|
| **DorsAttn** | age | 3.75 | <0.001 | 0.0012 | ** |
| | sex | -2.91 | 0.0050 | 0.016 | * |
| **VentAttn** | age | 2.67 | 0.0096 | 0.012 | * |
| | sex | 2.67 | 0.0096 | 0.024 | * |
| **Limbic** | age | 2.050 | 0.044 | 0.044 | * |
| | sex | -2.26 | 0.027 | 0.037 | * |
| **Control** | age | 4.87 | <0.001 | <0.001 | *** |
| | sex | -2.26 | 0.027 | 0.039 | * |
| **Default** | age | 2.82 | 0.0064 | 0.0091 | ** |
| | sex | -3.11 | 0.0027 | 0.016 | * |
| **Subcortical** | age | 2.86 | 0.0057 | 0.0091 | ** |
| | sex | -1.48 | 0.14 | 0.16 | ns |
| **Cerebellum** | age | 3.84 | <0.001 | 0.0012 | ** |
| | sex | -1.41 | 0.16 | 0.16 | ns |
| **Spinal cord** | age | 2.93 | 0.0046 | 0.0091 | ** |
| | sex | -1.89 | 0.062 | 0.078 | ns |

**Table S9 | Age and sex effects on inter-network functional connectivity.** The age-related effect on inter-network was obtained for each brain network and the spinal cord using an ordinary least squares (OLS) model including age and sex as fixed effects. The degree of freedom was always 65 (*i.e.,* 67 individuals – 2 fixed factors) and the t, p-values and FDR-corrected p-values are reported for each fixed effect. For the sex effect, Female is the reference level. The coefficient (and its t-value) represents the extent to which the mean outcome for Males differs from that for Females; thus, a positive value indicates higher values in Males *compared to* Females, and *vice versa*. The last column represents the significance: ns: not significant, . : significant before FDR but not after, *: $p_{FDR} < 0.05$, **: $p_{FDR} < 0.01$; ***: $p_{FDR} < 0.001$.

| Networks | Fixed effect | t-value | p-value | p-value FDR | Significance |
|---|---|---|---|---|---|
| **Visual** | age | 4.36 | <0.001 | <0.001 | *** |
| | sex | -2.24 | 0.029 | 0.10 | ns |
| **SomMot** | age | 0.18 | 0.85 | 0.85 | ns |
| | sex | 0.41 | 0.68 | 0.79 | ns |
| **DorsAttn** | age | 2.80 | 0.0067 | 0.016 | * |
| | sex | -2.08 | 0.041 | 0.10 | . |
| **VentAttn** | age | 1.97 | 0.052 | 0.10 | ns |
| | sex | 0.040 | 0.97 | 0.97 | ns |
| **Limbic** | age | 1.63 | 0.11 | 0.13 | ns |
| | sex | -0.55 | 0.59 | 0.79 | ns |
| **Control** | age | 1.79 | 0.077 | 0.13 | ns |
| | sex | 0.36 | 0.71 | 0.79 | ns |
| **Default** | age | 4.51 | <0.001 | <0.001 | *** |
| | | 0.76 | | | ns |

| | | | | | |
|---|---|---|---|---|---|
| | sex | | 0.45 | 0.75 | |
| **Subcortical** | age | 1.72 | 0.090 | 0.13 | ns |
| | sex | 2.16 | 0.035 | 0.10 | . |
| **Cerebellum** | age | 5.11 | <0.001 | <0.001 | *** |
| | sex | -1.60 | 0.11 | 0.22 | ns |
| **Spinal cord** | age | 0.85 | 0.39 | 0.44 | ** |
| | sex | -3.07 | 0.0030 | 0.030 | * |

**Table S10 | Age and sex effects on intra-network functional connectivity.** The age-related effect on intra-network was obtained for each brain network and the spinal cord using an ordinary least squares (OLS) model including age and sex as fixed effects. The degree of freedom was always 65 (*i.e.,* 67 individuals – 2 fixed factors) and the t, p-values and FDR-corrected p-values are reported for each fixed effect. For the sex effect, Female is the reference level. The coefficient (and its t-value) represents the extent to which the mean outcome for Males differs from that for Females; thus, a positive value indicates higher values in Males *compared to* Females, and *vice versa*. The last column represents the significance: ns: not significant, . : significant before FDR but not after, *: $p_{FDR} < 0.05$, **: $p_{FDR} < 0.01$; ***: $p_{FDR} < 0.001$.

| Feature | Rois | Fixed effect | t-value | p-value | p-value FDR | Signif. |
|---|---|---|---|---|---|---|
| | **Visual** | age | 1.75 | 0.081 | 0.30 | ns |
| | | sex | 0.23 | 0.82 | 0.93 | ns |
| | **SomMot** | age | 1.88 | 0.060 | 0.25 | ns |
| | | sex | 1.38 | 0.17 | 0.33 | ns |
| | **DorsAttn** | age | -0.92 | 0.35 | 0.70 | ns |
| | | sex | 0.74 | 0.46 | 0.63 | ns |
| | **VentAttn** | age | 3.14 | 0.0017 | 0.027 | * |
| | | sex | 2.29 | 0.022 | 0.097 | . |
| | **Limbic** | age | 3.27 | 0.0011 | 0.022 | * |
| | | sex | 0.86 | 0.39 | 0.58 | ns |
| **SD** | **Control** | age | 1.91 | 0.056 | 0.25 | ns |
| | | sex | 1.36 | 0.17 | 0.33 | ns |
| | **Default** | age | 3.35 | <0.001 | 0.019 | * |
| | | sex | 1.11 | 0.27 | 0.46 | ns |
| | **Subcortical** | age | 3.84 | <0.001 | 0.0086 | ** |
| | | sex | 2.48 | 0.013 | 0.069 | . |
| | **Cerebellum** | age | 3.64 | <0.001 | 0.0095 | ** |
| | | sex | 1.37 | 0.17 | 0.32 | ns |
| | **Spinal cord dorsal** | age | 3.74 | <0.001 | 0.0095 | ** |
| | | sex | 1.70 | 0.087 | 0.21 | ns |
| | **Spinal cord ventral** | age | 1.58 | 0.11 | 0.39 | ns |
| | | sex | 0.17 | 0.86 | 0.93 | ns |

| | | | | | | |
|---|---|---|---|---|---|---|
| | **Visual** | age | 2.63 | 0.0084 | 0.080 | . |
| | | sex | 1.95 | 0.050 | 0.16 | ns |
| | **SomMot** | age | 3.66 | <0.001 | 0.0096 | ** |
| | | sex | 2.29 | 0.022 | 0.097 | . |
| | **DorsAttn** | age | 1.96 | 0.050 | 0.24 | ns |
| | | sex | 1.71 | 0.086 | 0.21 | ns |
| | **VentAttn** | age | 3.42 | <0.001 | 0.016 | * |
| | | sex | 2.78 | 0.0054 | 0.051 | . |
| | **Limbic** | age | 3.84 | <0.001 | 0.0086 | ** |
| | | sex | 1.94 | 0.052 | 0.16 | ns |
| | **Control** | age | 3.52 | <0.001 | 0.012 | * |
| | | sex | 2.07 | 0.038 | 0.13 | . |
| | **Default** | age | 4.24 | <0.001 | 0.0045 | ** |
| **ALFF** | | sex | 2.13 | 0.033 | 0.12 | . |
| | **Subcortical** | age | 2.19 | 0.028 | 0.18 | . |
| | | sex | 2.58 | 0.0098 | 0.061 | . |
| | **Cerebellum** | age | 2.93 | 0.003 | 0.044 | * |
| | | sex | 2.67 | 0.007 | 0.058 | . |
| | **Spinal cord dorsal** | age | 2.84 | 0.0044 | 0.048 | * |
| | | sex | 4.33 | <0.001 | 0.0031 | ** |
| | **Spinal cord ventral** | age | 0.82 | 0.41 | 0.73 | ns |
| | | sex | 3.012 | 0.0026 | 0.051 | . |

**Table S11 | Age and sex effects on functional dynamic profiles in both brain and spinal cord regions.** The age-related effect was obtained for each feature and four regions using an ordinary least squares (OLS) model including age and sex as fixed effects. We report the fixed effects of the model when the age effect was significant before FDR correction for at least one region. The degree of freedom was always 65 (*i.e.,* 67 individuals – 2 fixed factors) and the t, p-values and FDR-corrected p-values are reported for each fixed effect. For the sex effect, Female is the reference level. The coefficient (and its t-value) represents the extent to which the mean outcome for Males differs from that for Females; thus, a positive value indicates higher values in Males *compared to* Females, and *vice versa*. The last column represents the significance: ns: not siginifcant, . : significant before FDR but not after, *: $p_{FDR} < 0.05$, **: $p_{FDR} < 0.01$; ***: $p_{FDR} < 0.001$. DL: dorsal left, DR: dorsal right, VL: ventral left, VR: ventral right, SD: Standard deviation, ALFF: amplitude of low frequency fluctuation, ACF: autocorrelation function.
